# Analysis of long and short enhancers in melanoma cell states

**DOI:** 10.1101/2021.07.27.453936

**Authors:** David Mauduit, Liesbeth Minnoye, Ibrahim Ihsan Taskiran, Maxime de Waegeneer, Valerie Christiaens, Gert Hulselmans, Jonas Demeulemeester, Jasper Wouters, Stein Aerts

## Abstract

Understanding how enhancers drive cell type specificity and efficiently identifying them is essential for the development of innovative therapeutic strategies. In melanoma, the melanocytic (MEL) and the mesenchymal-like (MES) states present themselves with different responses to therapy, making the identification of specific enhancers highly relevant. Using massively parallel reporter assay (MPRA) in a panel of patient-derived melanoma lines (MM lines), we set to identify and decipher melanoma enhancers by first focusing on regions with state specific H3K27 acetylation close to differentially expressed genes. A more in-depth evaluation of those regions was then pursued by investigating the activity of ATAC-seq peaks found therein along with a full tiling of the acetylated regions with 190 bp sequences. Activity was observed in more than 60% of the selected regions and we were able to precisely locate the active regions within ATAC-seq peaks. Comparison of sequence content with activity, using the deep learning model DeepMEL2, revealed that AP-1 alone is responsible for the MES enhancer activity, while SOX and MITF both influence MEL enhancer activity with SOX being required to achieve high levels of activity. Overall, our MPRA assays shed light on the relationship between long and short sequences in terms of their sequence content, enhancer activity, and specificity as reporters across melanoma cell states.

## Introduction

Enhancers are crucial regulatory regions in the genome that control cell type-specific gene expression. Identifying enhancers helps to better understand cell identity and is key to develop therapies targeting a singular relevant cell type in a disease. To date, the accurate prediction of enhancer location and activity in a given cell type remains a challenge. Both the presence and clustering of transcription factor binding sites (TFBSs) are good predictors of enhancer activity (Gasperini et al., 2020; King et al., 2020). Yet, such an approach requires prior knowledge of the cis-regulatory grammar in the studied cell types as only a small proportion of the TFBSs found in the genome are bound by the corresponding transcription factor (TF) (Yáñez-Cuna et al., 2012). Another strategy to identify candidate enhancers is to use active enhancer marks such as H3K27ac and chromatin accessibility (Gray et al., 2017; Minnoye et al., 2021; Rada-Iglesias et al., 2011). The most successful studies, combining this approach with transcriptome data, generated libraries with up to 60% of active enhancers in the target cell type (Gorkin et al., 2020; Graybuck et al., 2021). Massively parallel reporter assays (MPRA) have been developed to screen the activity of thousands of sequences simultaneously (Inoue and Ahituv, 2015; Melnikov et al., 2012; White et al., 2013). However, limitations of sequence synthesis constrain one to choose either a large number of short sequences (e.g., thousands of sequences of 150-250 bp) or a small number of longer sequences (e.g., dozens of sequences of 500-1000 bp)(Inoue and Ahituv, 2015). This issue, combined with the difficulty to identify putative enhancers, leads to a low rate of active enhancers in MPRAs.

Here, we study enhancer location, specificity and regulatory grammar in melanoma, using a variety of MPRA strategies. Melanoma exhibits pronounced heterogeneity within and between patients (Grzywa et al., 2017). Two main subtypes or cell states are discernable, melanocytic (MEL) and mesenchymal-like (MES) (Hoek et al., 2008, 2006; Verfaillie et al., 2015), as well as more recently identified variants of the MEL state, such as the neural-crest like and intermediate states (Rambow et al., 2018; Tsoi et al., 2018; Wouters et al., 2020). MEL and MES subtypes display distinct epigenomic and transcriptomic profiles resulting in divergent phenotypes (e.g., migration; Wouters et al., 2020), and different responses to therapy (Verfaillie et al., 2015). Thus, the identification of subtype-specific enhancers may be relevant for therapy, where it could improve safety and efficiency by narrowing down the effect of the treatment to a specific population. Comparisons between MEL and MES yield thousands of regions with differential acetylation (H3K27ac) and accessibility. However, it remains unclear which of these subtype-specific regions function as active enhancers and which TFs are responsible for their activity.

In this study, we analyzed MEL-and MES-specific regions identified based on differential H3K27ac chromatin immunoprecipitation sequencing (ChIP-seq) signal nearby differentially expressed genes. We designed MPRA experiments to test those regions at three different levels in a panel of patient-derived malignant melanoma (MM) lines. Our results precisely locate the origins of enhancer activity within the larger H3K27ac domains. In addition, we can accurately predict their subtype specificity, and ultimately identify a set of rules governing MEL and MES enhancer activity. Furthermore, we show that a melanoma deep learning model (DeepMEL2; Atak et al., 2021) trained on ATAC-seq data pinpoints which TFBSs drive enhancer activity and specificity.

## Results

### Design of MPRA libraries based on H3K27ac, ATAC-seq, and synthetic sequences

H3K27ac ChIP-seq peaks are often used for the selection of candidate enhancers (Creyghton et al., 2010; Fox et al., 2020; Fu et al., 2018). However, such peaks can encompass large genomic regions, often 2-3 kb long, while enhancers are usually only a few hundred base pairs in size (Gasperini et al., 2020; Li and Wunderlich, 2017). To investigate the relationship between H3K27ac signal, chromatin accessibility peaks, and enhancer activity, we designed MPRA libraries at three different levels: 1.2 to 2.9kb sized H3K27ac ChIP-seq peaks, 501bp sized ATAC-seq peaks that fall within the H3K27ac regions, and 190 bp subsequences tiling the entire H3K27ac regions.

We designed the H3K27ac ChIP-seq based library (H3K27ac library) by selecting regions that are specifically acetylated in either the MEL or the MES melanoma cell state and located around differentially expressed genes in a panel of 12 melanoma lines: 3 MES lines (MM029, MM047 and MM099) and 9 MEL lines that cover a spectrum from pure-melanocytic to intermediate melanoma (MM001, MM011, MM031, MM034, MM057, MM074, MM087, MM118, SKMEL5) (See Methods, Fig 1a.; Minnoye et al., 2020). A special consideration was given to regions overlapping with ChIP-seq peaks for SOX10 and MITF (for MEL regions) or AP-1 (JUN and JUNB; for MES regions), known regulators of each state. A total of 35 MES-and 18 MEL-specific regions, with an average size of 1,987 bp, were amplified from genomic DNA (Supplementary Table 1). The H3K27ac ChIP-seq signal across the selected regions displays a good correlation between cell lines of the same subtype and a negative correlation between cell lines of a different subtype (Supplementary Fig 1a.). This correlation is also observed in the ATAC-seq signal and the target gene expression (Supplementary Fig 1b.,c.,g.). We created two vector libraries based on the specific CHEQ-seq vector backbone (Verfaillie et al., 2016), by cloning the sequences upstream (5’ position) or downstream (intron position) of a minimal promoter (SCP1, see Methods, Fig 1b. left panel).

**Figure 1:**
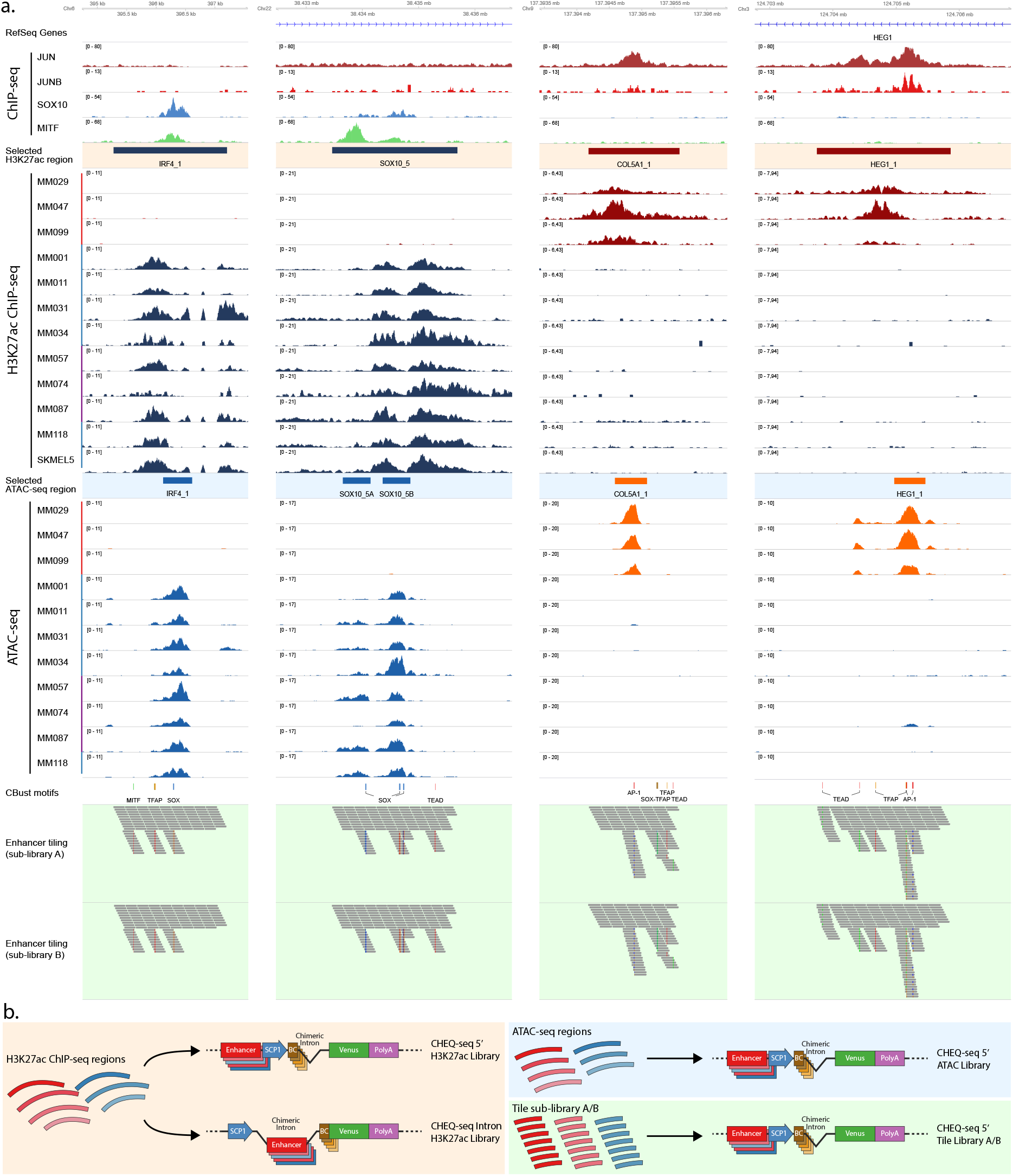
**a.**, Cell state-specific regions were selected based on H3K27ac ChIP-seq signal from a panel of melanoma cell lines containing both MES (red bar) and MEL lines (pure MEL: blue bar and intermediate MEL: purple bar). ATAC-seq data from the same lines was used to identify accessibility peaks within these regions. Finally, the regions were tiled with 190 bp tiles with a shift of 20 bp (sub-library A). Sub-library B was generated by shifting all tiles 10 bp downstream. CBust was used to identify TF motifs and new tiles were generated with mutated motifs. **b.**, Reporter vector configurations used for the evaluation of the H3K27ac enhancers (left panel), the ATAC-seq enhancers (top right panel) and the enhancer tiling (bottom right panel). SCP1: Super Core Promoter 1, BC: Barcode.

Next, we designed a second library consisting of the ATAC-seq peaks contained within the H3K27ac library regions (Fig 1a.). In some cases, two peaks were selected within the same region. Each sequence was defined by taking the summit of the ATAC-seq peak and extending 250 bp on each side, resulting in a 501 bp candidate sequence. 28 MES-and 18 MEL-specific ATAC-seq peaks were selected. The correlation of the H3K27ac ChIP-seq, ATAC-seq and RNA-seq signal between the different cell lines remains the same as for the H3K27ac library (Supplementary Fig 1d.-f.). We cloned the regions in the CHEQ-seq vector, upstream of the SCP1 promoter (Fig 1b. upper right panel). In addition, we cloned the same set of sequences in the STARR-seq vector (an alternative MPRA vector) to assess assay-related variability (Muerdter et al., 2017).

Finally, to locate the precise origin of enhancer activity, we generated two tiling libraries (A and B, see Methods), encompassing the entire H3K27ac regions. The tiles are 190 bp long with a 20 bp shift between consecutive tiles. The tiling for library A starts at nucleotide position 1 of the H3K27ac ChIP-seq regions while library B starts at position 11, resulting in a final tiling resolution of 10 bp when both sub-libraries are taken into account.

In order to probe the effect of mutations within putative TFBSs, we used Cluster-Buster (cbust; Frith et al., 2003) with position weight matrices for binding sites of key regulators of MEL (SOX10, MITF, TFAP2A) and MES (AP-1 and TEAD) (Wouters et al., 2020) to identify TFBSs present in the sequences of the H3K27ac library. We generated tiles with mutated versions of these motifs (See Methods). For each sub-library, 800 shuffled tiles were generated as negative controls, resulting in a total of 7,412 and 7,356 tiles for sub-library A and B, respectively (Fig 1a.). Each sub-library is separately cloned upstream of the SCP1 promoter in the CHEQ-seq vector and is transfected individually (Fig 1b. lower right panel).

### Most MEL-specific acetylated regions harbour enhancer activity in MEL lines

We first transfected all MPRA libraries in the most melanocytic (MEL) line, MM001 (Minnoye et al., 2020; Wouters et al., 2020). Of the MEL-specific H3K27ac regions, 75% (14/18) display significant enhancer activity (Benjamini–Hochberg adjusted p-values < 0.05, see Methods) in MM001 with a mean log2 fold change (FC) of 0.23, compared to 26% (8/32) for the MES-specific regions with a mean log2 FC of −1.21 (Fig 2a.). The activities are consistent across the two library designs (enhancers cloned into the intron or upstream of the TSS; Supplementary Fig 2a.-b.). The library where only the ATAC-seq peaks were cloned recapitulates these activities, suggesting that the enhancer activity is contained within the sequence of the ATAC-seq peak (Fig 2b., Supplementary Fig 2c.-e.). Interestingly, in the five MEL-specific H3K27ac regions where two ATAC-seq peaks were assessed, only one of the two recapitulates the activity of the encompassing region (Supplementary Fig 3). This was independently confirmed using the STARR-seq MPRA (Supplementary Fig 2c.).

**Figure 2:**
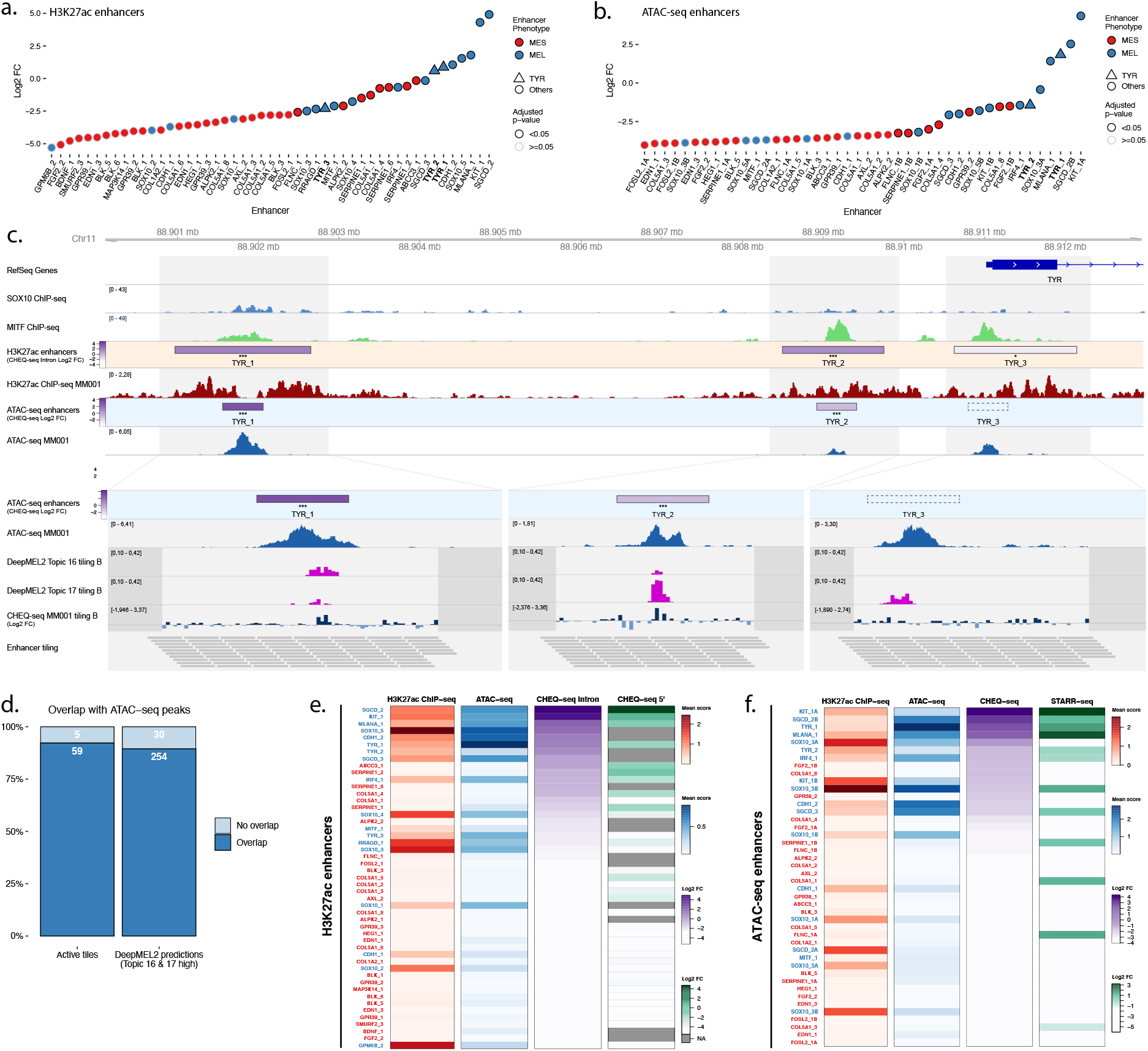
Enhancer activity in MM001. **a.**-**b.**, Enhancer activity profile for the CHEQ-seq intron H3K27ac library (**a.**) and CHEQ-seq ATAC-seq library (**b.**). Enhancer regions displayed in panel **c.** have their name indicated in bold and their value is displayed with a triangle. **c.**, Enhancer activity of regions selected around the TYR genes. SOX10 and MITF ChIP-seq as well as H3K27ac ChIP-seq and ATAC-seq for MM001 are displayed and, in the zoomed-in regions (light grey areas), DeepMEL2 predictions and CHEQ-seq values of the enhancer tiling B library are represented. Dark grey areas are regions not covered by a tile. CHEQ-seq activity is visible in the ‘H3K27ac enhancers’ and ‘ATAC-seq enhancers’ tracks. Benjamini–Hochberg adjusted p-values: * < 0.05; *** < 0.001. Dashed box: region not recovered following DNA synthesis, cloning or MPRA assay. **d.**, Percentage of overlap between active tiles and ATAC-seq peaks (left) and high DeepMEL2 predictions with ATAC-seq peaks (right). **e.**-**f.**, Heatmaps of H3K27ac library (**e.**) and ATAC-seq library (**f.**), displaying H3K27ac ChIP-seq signal, ATAC-seq signal and enhancer activity in MM001 ordered by MPRA values. Only MPRA values of significantly active enhancers are displayed.

Next, we examined the activity of all 190 bp sequences tiled along the entire H3K27ac regions. We confirmed that the majority of active tiles (92.2%) are located within an ATAC-seq peak (Fig 2c.-d.; Supplementary Fig 3) and identified active tiles in 7 out of the 10 most active MEL ATAC-seq based enhancers. Short 190 bp regions can thus often recapitulate the enhancer activity of the larger encompassing region. When two or more consecutive tiles, that are shifted 20bp, are active, the enhancer may be contained in an even smaller sequence or the activity is coming from independently active elements close to one another.

We recently trained a deep learning model on cis-regulatory topics from 30 melanoma lines, called DeepMEL2, that accurately predicts the accessibility and activity of a sequence in the different melanoma subtypes (Atak et al., 2021). Each topic used to train the model regrouped accessible regions found in one cell line, in a specific subtype or in all cell lines. Two topics are associated with the MEL subtype, topic 16 and 17, mostly focused on SOX and MITF motifs respectively. We scored our 190 bp tiles with DeepMEL2 and found high MEL prediction scores (>0.10, see Methods) specifically within ATAC-seq peaks (Fig 2d.). Of those top DeepMEL2 predictions for MEL specific topics, 11% (Topic 16) and 17.3% (Topic 17) are active tiles in the MPRA (with 0.25/0.375 recall and 0.11/0.173 precision for topic 16 and 17 respectively). These low precision values may be explained by the fact that the DeepMEL2 model was trained on ATAC-seq data, thus yielding high prediction scores within ATAC-seq peaks, yet not all of these show positive MPRA activity.

In some cases, we identified multiple acetylated regions near one gene. For the tyrosinase (*TYR*) gene, expressed specifically in MEL lines (Supplementary Fig 1g.), three regions were selected as MEL-specific and tested at the acetylation, accessibility and tiling levels (Fig 2c.). TYR_1 and TYR_2 regions display high reporter activity which is subsequently found in the selected ATAC-seq peak. Activity is further found in tiles at the same location as the DeepMEL2 predictions (Fig 2c.). The TYR_3 region, at the gene’s promoter site, has a low activity that is also not found when tiling the enhancer, despite the DeepMEL2 predictions. Those findings suggest that TYR expression in MEL lines is largely dependent on the activity of distal enhancers.

Some other enhancers that are active in the H3K27ac and ATAC-seq libraries, are not recapitulated in the tile library (e.g. SOX10_3 Supplementary Fig 3c.). This can be due to technical reasons, such as the small size of the tiles. Nevertheless, from the combination of the ATAC-seq and enhancer tiling MPRAs we can conclude that not all subtype-specific ATAC-seq peaks function as a standalone enhancer.

We finally compared signals of H3K27ac ChIP-seq mean score, ATAC-seq mean score and their corresponding MPRA signals (Fig 2e.-f.). With the H3K27ac library (Fig 2e.), the acetylation signal measured over the selected regions correlates well with the accessibility signal (Pearson’s correlation r = 0.77) indicating that ATAC-seq peaks are found within the selected acetylated regions and maintain the same differential signal. Active enhancers are found in the majority (14/18) of the MEL-specific acetylation regions with ATAC-seq signal. This trend is also visible in the ATAC-seq based library, with most of the active enhancers detected in ATAC-seq peaks (13/19) (Fig 2f.). However, the moderate correlation between ATAC-seq signal and CHEQ-seq activity in the corresponding library (Spearman’s rho = 0.48) also indicates that the peak mean signal is not a good predictor of the activity level of an enhancer. In part, this may be due to confounding of ATAC and H3K27ac read depth by genomic copy number aberrations. Also, the activity displayed by some MES-specific regions lacking ATAC-seq signal in MM001 suggests that closed regions in the genome can still harbour activity in an episomal MPRA assay.

In conclusion, our enhancer selection resulted in a high rate of active enhancers in MM001 and the design of our MPRA libraries allowed us to precisely pinpoint the origins of, at least part of, the enhancer activity.

### MES-specific H3K27ac/ATAC regions are active in MES lines

Next, we transfected all libraries in the MES line MM029. The activity profiles in this line show that, as expected, the majority of MES enhancers display activity at both the H3K27ac and ATAC-seq level (Fig 3a.,b.; Supplementary Fig 4a.-e.). In regions with two selected ATAC-seq peaks, both MPRA approaches agree, once again, on which peak is driving activity (Fig 3b.,d.; Supplementary Fig 4c.; Supplementary Fig 5).

**Figure 3:**
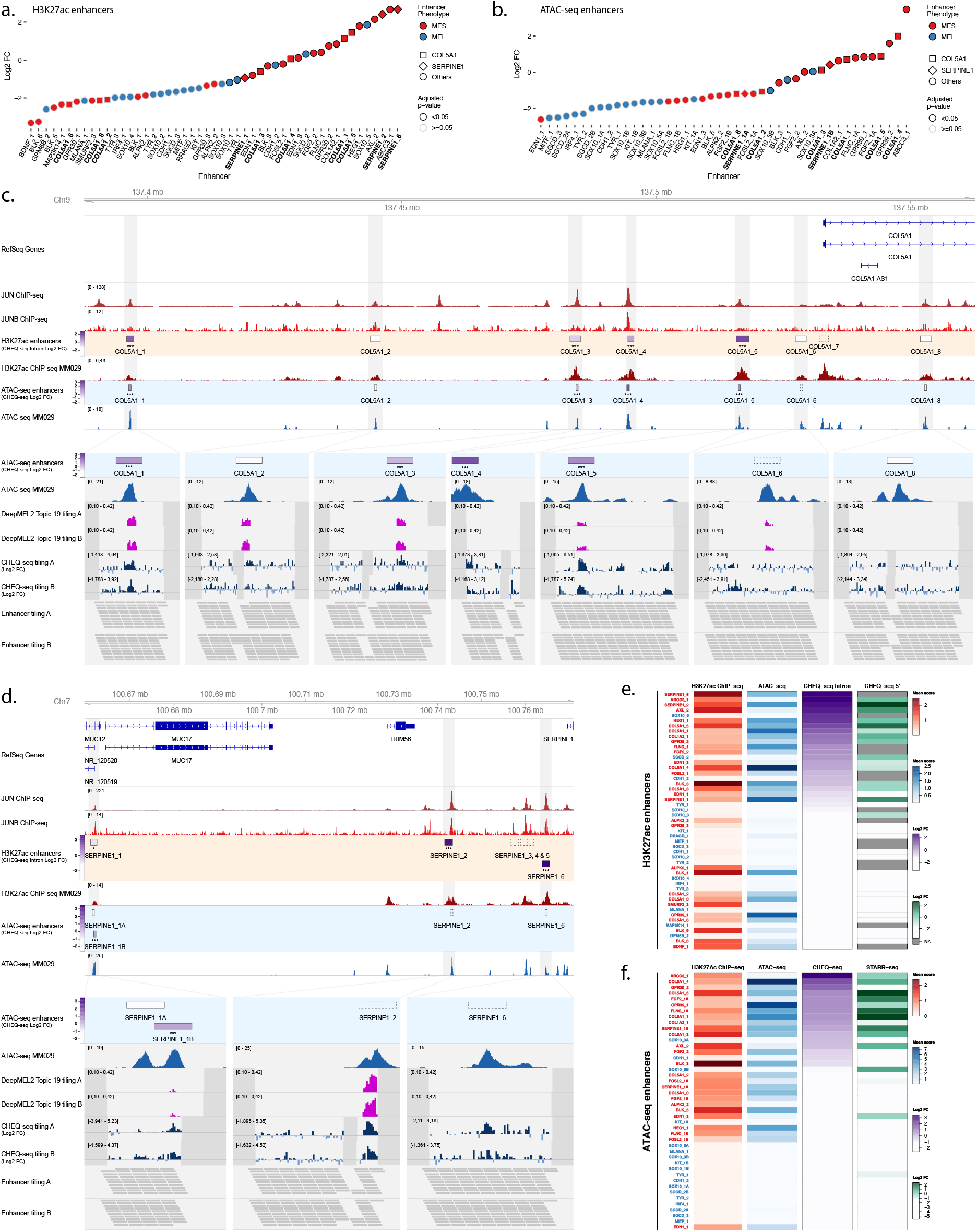
Enhancer activity in MM029. Enhancer activity profile for the CHEQ-seq intron H3K27ac library (**a.**) and CHEQ-seq ATAC-seq library (**b.**). Enhancer regions displayed in panel **c.** and **d.** have their name indicated in bold and their value is displayed with a different shape. Enhancer activity of regions selected around the COL5A1 (**c.**) and SERPINE1 (**d.**) genes. JUN and JUNB ChIP-seq and H3K27ac ChIP-seq and ATAC-seq for MM029 are displayed and, in the zoomed-in regions (light grey areas), DeepMEL2 predictions and CHEQ-seq values of the enhancer tiling are represented. Dark grey areas are regions not covered by the tiling library. CHEQ-seq activity is visible in the **‘**H3K27ac enhancers’ and ‘ATAC-seq enhancers’ tracks. Benjamini–Hochberg adjusted p-values: * < 0.05; *** < 0.001. Dashed boxes: regions not recovered following DNA synthesis, cloning or MPRA assay. Heatmaps of H3K27ac library (**f.**) and ATAC-seq library (**g.**), displaying H3K27ac ChIP-seq signal, ATAC-seq signal and enhancer activity in MM001 ordered by MPRA values. Only MPRA values of significantly active enhancers are displayed.

Multiple regions around the Collagen Type V Alpha 1 Chain (*COL5A1*) gene were found specifically acetylated in MES lines, up to 100kb upstream of the TSS, and we included a total of seven into the library (Fig 3c.). Four regions showed significant activity in MM029 (Fig 3c.). This activity is further confirmed in the ATAC-seq and tiling libraries: the ATAC-seq peaks within the four active H3K27ac regions are all active, while the ATAC-seq peaks within the three negative H3K27ac regions are all negative. In the COL5A1_5 region, tiling suggests the presence of three distinct enhancers, including two that are located within small ATAC-seq peaks that were not selected for the ATAC-seq based library.

DeepMEL2 is trained on both MEL and MES accessible regions, and topic 19 has been shown to be the best performing MES topic (Atak et al., 2021). In our data, topic 19 high predictions often (18.1%) overlap with active tiles in MM029 (0.181 precision, 0.537 recall, Fig 3c.-d.). Because several other cis-regulatory topics contribute to the prediction of the MES subtype, some tiles do not display a high prediction score for topic 19 despite their activity and can be better explained by other topics.

The small shift between each tile and the use of two overlapping libraries provides a high number of measurements throughout the regions which allows for the more accurate detection of lowly active enhancers. Such enhancers are found in the SERPINE1_1 region, where the SERPINE1_1A ATAC-seq peak is inactive in the CHEQ-seq and STARR-seq assays (Fig 3d., Supplementary Fig 4c.) but the tiling assay shows robust enhancer activity. Of the two ATAC-seq peaks in the FOSL2_1 region, neither one recapitulates the activity of the acetylated region. In contrast, the tiling assay reveals clear enhancer activity in both peaks (Supplementary Fig 5b.).

Similar conclusions to those above for MEL enhancers, can now be drawn for MES regions regarding the relationship between H3K27ac, ATAC-seq signal and enhancer activity (Fig 3e.-f.). ATAC-seq based and tiling CHEQ-seq assays show that most active enhancers in the H3K27Ac regions reside within ATAC-seq peaks (151/164 active tiles are found in ATAC-seq peaks; 92.1%). Irrespective, ATAC-seq peak mean signal remains a poor predictor of the level of enhancer activity, at least as read out by CHEQ-seq. Moreover, the presence of a differentially accessible ATAC-seq peak does not guarantee enhancer function. Indeed, of 26 differentially accessible peaks, only 14 show demonstrable activity (54%).

In conclusion, MPRA assays performed in MM001 and MM029 have shown a high success rate of MEL and MES selected regions to display activity specifically in their corresponding cell state. However, MM001 and MM029 lie at the extremes of the MEL-MES spectrum. To further investigate how the activity of the selected regions scales along this axis, we studied them in five additional melanoma cell lines, representing more intermediate or transitory melanoma states (Tsoi et al., 2018; Wouters et al., 2020).

### MES enhancers show lower but consistent activity in intermediate lines

To further study the behaviour of melanoma enhancers, we expanded our panel of cell lines to include two additional MES lines (MM047 and MM099) and three MEL-intermediate lines (MM057, MM074 and MM087). These three lines have high SOX10 and MITF expression (hallmarks of the MEL subtype), yet also show both marker expression and phenotypic characteristics typical for the MES subtype (e.g., AXL expression, TGFb1 signalling activity) (Tsoi et al., 2018). Furthermore, in contrast to MM001, these lines shift toward a MES subtype when SOX10 expression is lost (Wouters et al., 2020). MM057, MM074 and MM087 were part of the cell lines used for the selection of MEL-specific H3K27ac ChIP-seq regions. As such, they display an acetylation and accessibility profile as well as a transcriptional activity of the associated genes similar to MM001 (Supplementary Fig 1a.-c.). Based on those observations, even though phenotypically different, the MEL intermediate lines were expected to have an enhancer profile closely related to what we have observed with MM001.

The enhancer activity profile for the H3K27ac library obtained in intermediate lines correlates well with MM001, except for MM074 which moderately correlates with all lines (Fig 4a.). Interestingly, intermediate lines have the same proportion of active MEL enhancers as MM001 and the same proportion of active MES enhancers as the MES lines (Fig 4b.). When looking at the mean activity of each enhancer per cell line phenotype (Fig 4c.), we found a good correlation of MES region activity between all phenotypes (MES vs MM001 r = 0.64; MES vs Intermediate r = 0.89). The only difference resides in the strength of the enhancer activity where MM001 has very low activity and intermediate lines have moderate activity for MES regions.

**Figure 4:**
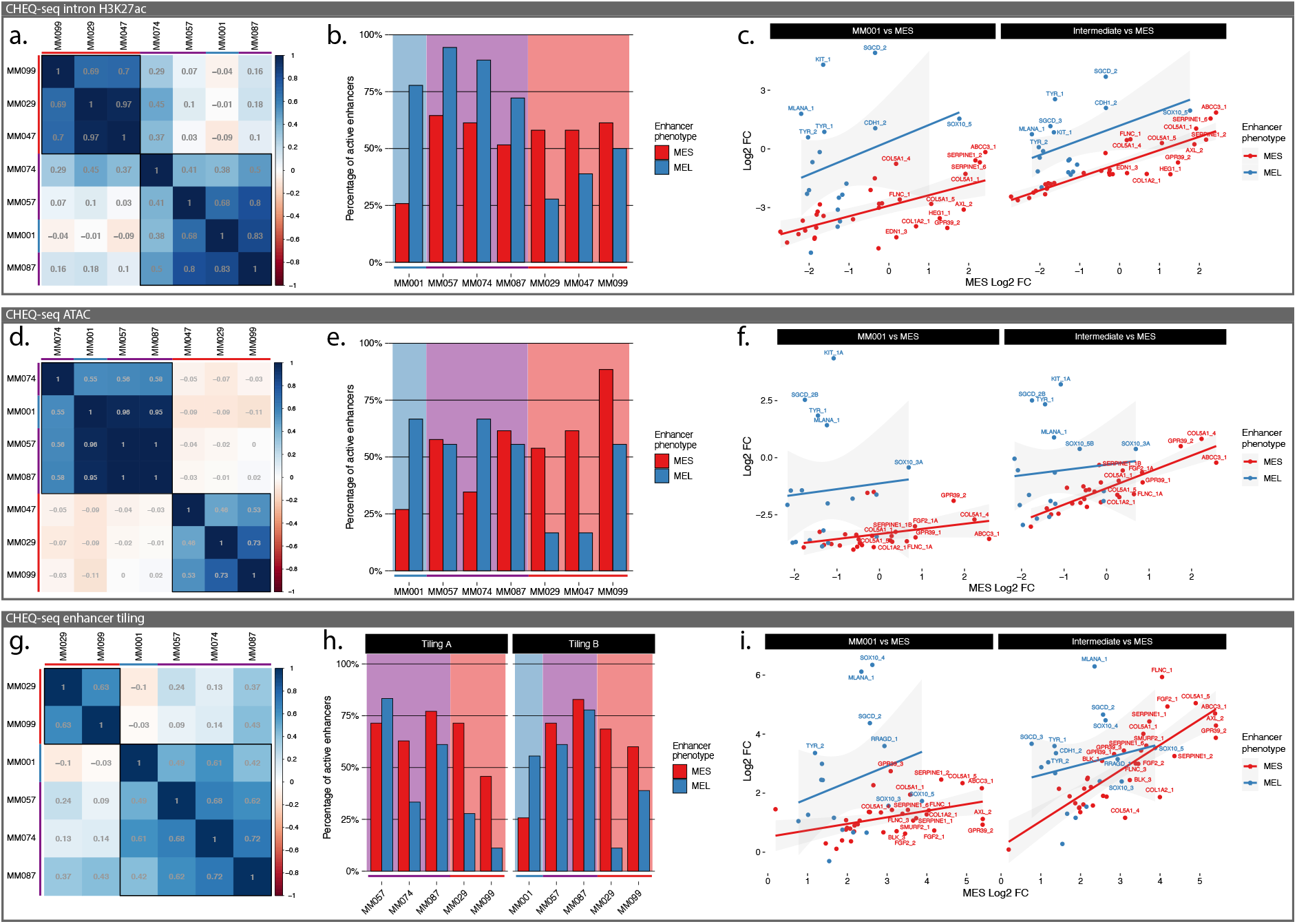
Specificity of MEL and MES enhancers in intermediate lines. **a., d., g.,** Pearson correlation coefficient table across 7 MM lines for CHEQ-seq intron H3K27ac library (**a.**), CHEQ-seq ATAC-seq library (**d.**) and combined CHEQ-seq enhancer tiling libraries (**g.**). **b.**, **e.**, **h.**, Percentage of active MEL and MES enhancers for each line (**b.**, H3K27ac library; **e.**, ATAC-seq library; **h.**, enhancer tiling libraries). Red, purple and blue bars next to cell line names and background indicate MES, Intermediate and MEL lines respectively. **c.**, **f.**, **i.**, Scatter plot of CHEQ-seq results for the intron H3K27ac library (**c.**), the ATAC-seq library (**f.**) and the combined CHEQ-seq enhancer tiling libraries (**i.**) activity with merged MES line values versus MM001 (left panel) or merged intermediate MEL lines (right panel).

The comparison of enhancer activity between subtypes highlights a particular case of the SOX10_5 region, a MEL-specific H3K27Ac region with enhancer activity in all cell lines (Fig 4c.). Based on the tiling profile of that region across all tested lines (Supplementary Fig 6a.), we can identify two active enhancers. One enhancer, located within the largest ATAC-seq peak, is MEL-specific and overlaps with the DeepMEL2 predictions for MEL accessibility (Green highlight, Supplementary Fig 6a.). The other one is located just downstream in a GC rich region and displays activity in all cell lines (Red highlight, Supplementary Fig 6b.). This profile explains why the SOX10_5B ATAC-seq region, which does not fully cover this GC rich region, is less active in MES lines (Fig 4f.).

With the ATAC-seq region library, the correlation between lines remains the same with a clear separation of the MEL and MES lines and the same pattern in the proportion of active enhancers (Fig 4d.-e.). The overall lower percentage of active MEL enhancers can be explained by the high proportion of regions with two peaks selected, where most of the time only one is active. As in the H3K27Ac library, the MES enhancer activity is higher in intermediate lines than in MM001 but much lower than in MES lines (Fig 4f.).

At the tiling level, intermediate lines show an increased MES enhancer activity to a level similar to the MES lines (Fig 4g.-i.). This results in a reduced correlation with MM001 despite a maintained strong activity of MEL enhancers. As observed in the specific part of the SOX10_5 region (Supplementary fig 6a.), most active MEL regions have a good specificity with no activity in MES lines. On the contrary, MES enhancers display a similar activity in MES lines and intermediate lines (Fig 4i.; Supplementary fig 6c.).

In summary, MEL enhancers display a consistent and specific activity in MEL lines while MES enhancers are not only active in MES lines but also in intermediate lines and to a lesser extent in MM001. This behaviour suggests that enhancer activity is driven by the expression of TFs responsible for the MEL and MES subtypes. SOX10, MITF and TFAP have been identified as drivers of the MEL state and AP-1 and TEAD of the MES state (Verfaillie et al., 2015). MM029 expresses MITF at a low level but it does not seem sufficient to drive the activity of MEL enhancers, suggesting that MITF alone cannot induce enhancer activity or that the expressed isoform in MM029 cannot bind the TFBS or activate the enhancer.

We will next investigate, for each state, which TFs are responsible for enhancer activity and whether a cis-regulatory grammar regulating the activity level can be observed.

### Key transcription factor binding sites explain melanoma enhancer activity

To better understand which TFs are important for MEL enhancer activity, we included tiles with MITF, SOX and TFAP binding site mutations in our tiling libraries. The comparison of wild type versus mutated tiles shows that TFAP does not affect the activity of the profiled enhancers in any of the MEL lines (Supplementary Fig 7). On the other hand, loss of SOX or MITF often negatively affects enhancer activity. KIT_1 provides a good example of both MITF and SOX motif contributions in MM001 (Fig 5a.). The tiling library confirms the location of enhancer activity to be in the KIT_1A region, where both motifs are found to influence the activity. Mutation of the SOX binding site abolishes tile activity. The same is observed when the MITF binding site is mutated, with the notable exception of one tile also containing the SOX motif. It is worth noting that despite KIT_1(A) being by far the most active region in both the H3K27ac (together with SGCD_2) and the ATAC-seq library, tiles with a higher activity have been found in 5 other MEL regions. This suggests that no individual tile recapitulates the enhancer activity coming from the whole accessible regions. The DeepMEL2 topic 16 and 17 prediction scores closely follow tile activity (FC), both highlighting the SOX and MITF motifs. Interestingly, the activity is not centred on the ATAC-seq peak summit. This suggests that, through training with ATAC-seq data, DeepMEL2 has identified TFBSs responsible for both accessibility *and* activity. Motif enrichment analysis on both the active and most accessible tiles in MEL lines identifies the E-box motif (MITF) as highly enriched in both active and accessible sequences while the SOX motif is only enriched in accessible tiles (Fig 4c.). On the other hand, AP-1 motifs are enriched only in active tiles. The discrepancy between the effect of some SOX mutations on activity and the absence of enrichment for the SOX motif in active tiles could be due to the low number of MEL enhancers tested (18).

**Figure 5:**
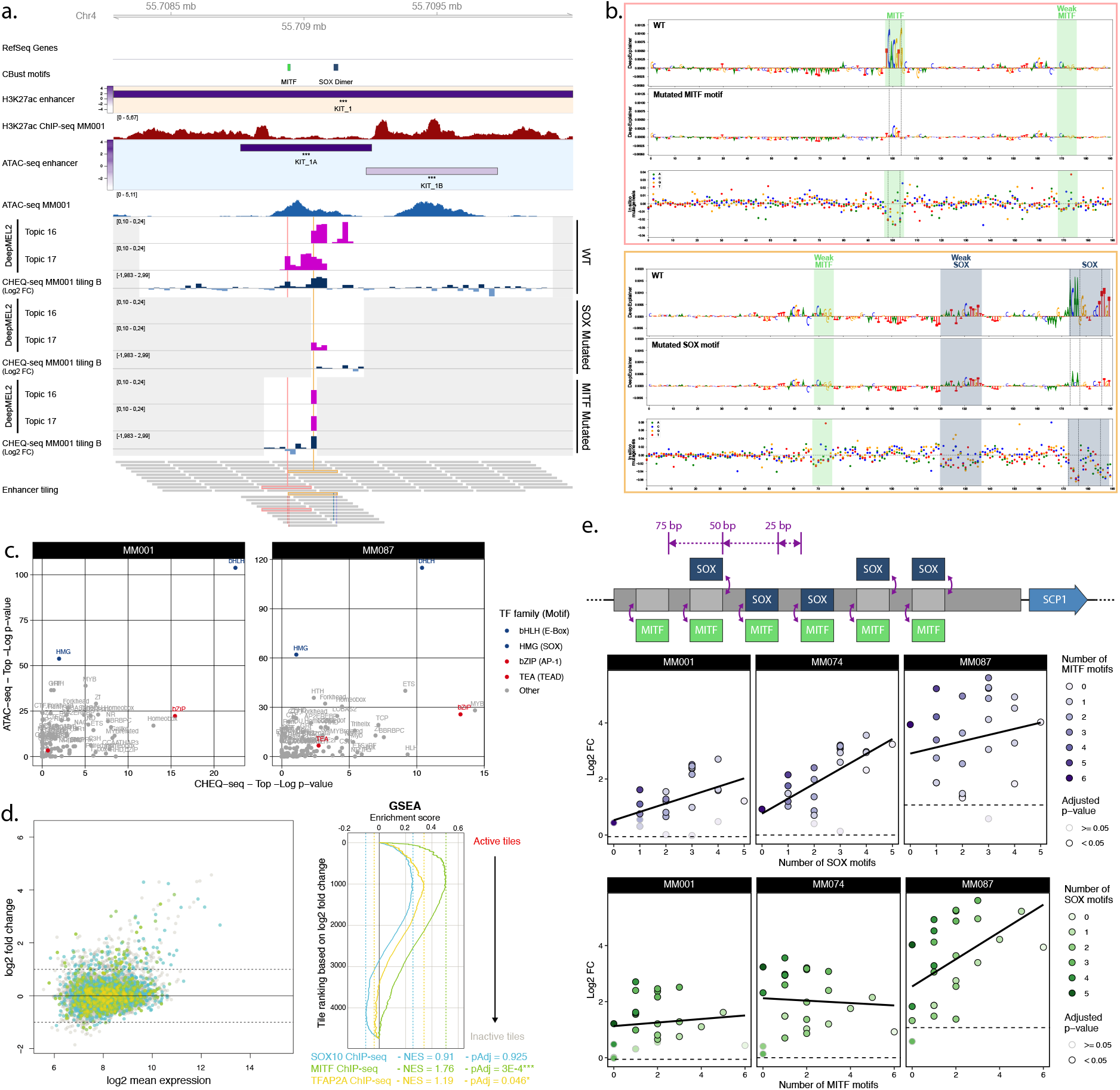
**a.**, KIT_1 region activity summary in MM001. The ‘CBust motifs’ track shows identified TFBSs for MITF and SOX. DeepMEL2 prediction scores for topic 16 (SOX) and 17 (MITF) and CHEQ-seq activity values are shown for the WT, SOX mutated and MITF mutated tiles. Grey areas are regions not covered by the tiling library. The ‘Enhancer tiling’ track represents the actual location of the tiles. **b.**, DeepExplainer profiles for the tiles highlighted in pink and yellow in the ‘Enhancer tiling’ track of panel **a.**. Each profile consists of the WT (top panel) and mutated (middle panel) tile nucleotide score and in silico mutagenesis score for the WT tile (bottom panel). Dashed lines indicate the location of mutated nucleotides. **c.**, Comparison of the most enriched motif families as identified by HOMER between ATAC-seq (top ¼ most accessible tiles vs rest) and CHEQ-seq tiling libraries (all active tiles vs all inactive tiles) for MM001 and MM087. **d.**, Tiles in the MA plot are coloured based on whether or not they overlap with SOX10 (blue), MITF (green) or TFAP2A (yellow) ChIP-seq peaks. GSEA of the enrichment of SOX10, MITF and TFAP2A ChIP-seq on the tiles ranked according to their activity (log2 FC). For each of the ChIP-seq peak sets, the negative enrichment score (NES) and Benjamini-Hochberg adjusted p-value (pAdj) are shown. **e.**, Top panel: cartoon of SOX and MITF motif combinations in a background sequence. Middle and bottom panels: CHEQ-seq activity of synthetic enhancers with background sequence 2 in MM001, MM074 and MM087 sorted by the number of SOX (middle panel) or MITF (bottom panel) motifs present in the sequence. Dashed line indicates the log2 FC value of the background sequence without any motif.

### SOX10 dependent ATAC-seq peaks with enhancer activity are enriched in MITF motifs

To study MEL enhancers in more detail, we designed a new library. A SOX10 knockdown (KD) with siRNA was performed on MM057 and MM087, shifting these lines to a MES phenotype, and was followed by ATAC-seq after 0, 24, 48 or 72 hours (Bravo González-Blas et al., 2019). After cisTopic analysis, we selected 1,461 ATAC-seq peaks where accessibility is lost upon KD and tiled them with 190 bp long sequences and 120 bp shifts, resulting in 6,696 individual sequences (Supplementary Fig 8a.-b.). We cloned this library in the CHEQ-seq vector and transfected it in MM087. Analysis of tile activity revealed that 15.1% of the selected ATAC-seq peaks exhibit enhancer function in MM087 (Supplementary Fig 8c.-d.). We performed Gene Set Enrichment Analysis (GSEA) of ChIP-seq of SOX10 and MITF in 501mel and TFAP2A in human primary melanocytes on the tiles ranked according to their activity (Fig 5d.). Only the MITF ChIP-seq signal was enriched in active tiles, indicating that SOX10-dependent regions containing MITF sites are preferentially active. Those observations were confirmed by differential motif discovery of active versus inactive tiles with the E-box motif (MITF) among the most strongly enriched motifs (Supplementary Fig 8e.).

### Synthetic SOX -MITF motifs combinations highlight MEL enhancer regulatory grammar

To further investigate a possible interaction between SOX10 and MITF and see if a MEL enhancer can be generated with only those 2 TFBSs, we designed sequences consisting of combinations of SOX dimer and MITF motifs spread over a 259 bp background sequence (Fig 5e. top panel). Twenty-four different combinations in 2 background sequences were generated, cloned in the CHEQ-seq vector and transfected in MM001, MM074 and MM087. The activity of the enhancer progressively increases with the number of SOX motifs (Fig 5e. middle panel). The presence of MITF motifs additionally increases enhancer activity but this differs from the effect of the SOX motifs (Fig 5e. bottom panel). We see a progressive increase of the activity based on the number of MITF binding sites only when there is also a SOX dimer motif present in the sequence. In the absence of a SOX motif, enhancer activity remains low in comparison with sequences containing at least one SOX motif, even though 6 MITF motifs are present. The presence of multiple SOX motifs greatly increases the enhancer activity, reducing the influence of MITF motifs (Fig 5e. bottom panel).

DeepMEL2 predictions on the synthetic sequences confirm these observations (Supplementary Fig 9a.-b.). Topic 16 predictions show the same constant increase based on the number of SOX binding sites in agreement with the measured activity. Topic 17 predictions also increase based on the number of MITF motifs when one SOX motif is present and are brought down to background level when SOX is absent. These results highlight a cis-regulatory grammar in MEL enhancers involving SOX10 and MITF. As previously described in enhancers regulating pluripotency in mouse embryonic stem cells, regardless of TFBSs positioning, the number of binding sites remains the predominant factor determining activity (Supplementary Fig 9c.; King et al., 2020).

### AP-1 sites alone can explain MES enhancer activity

Finally, we looked for the presence of AP-1 and TEAD binding sites in the previously tested acetylated regions and generated mutated versions of the enhancer tiles. Comparison of wild type versus TEAD or AP-1 mutant tiles shows a positive effect of AP-1 on activity in all cell lines (Supplementary Fig 10). On the other hand, TEAD motifs only show a limited effect in the MM029 tiling B sample and MM087 tiling A sample.

The COL5A1_5 region illustrates the influence of AP-1 (Fig 6a.). Within this region, three enhancers are found active in all cell lines bar MM001, which shows activity only for the second enhancer. The enhancer activity of the H3K27ac and ATAC based regions across the tested MM lines agrees with the tile activity where MM001 has no or low activity and intermediate and MES lines have a strong activity (Fig 6a.-b.).

**Figure 6:**
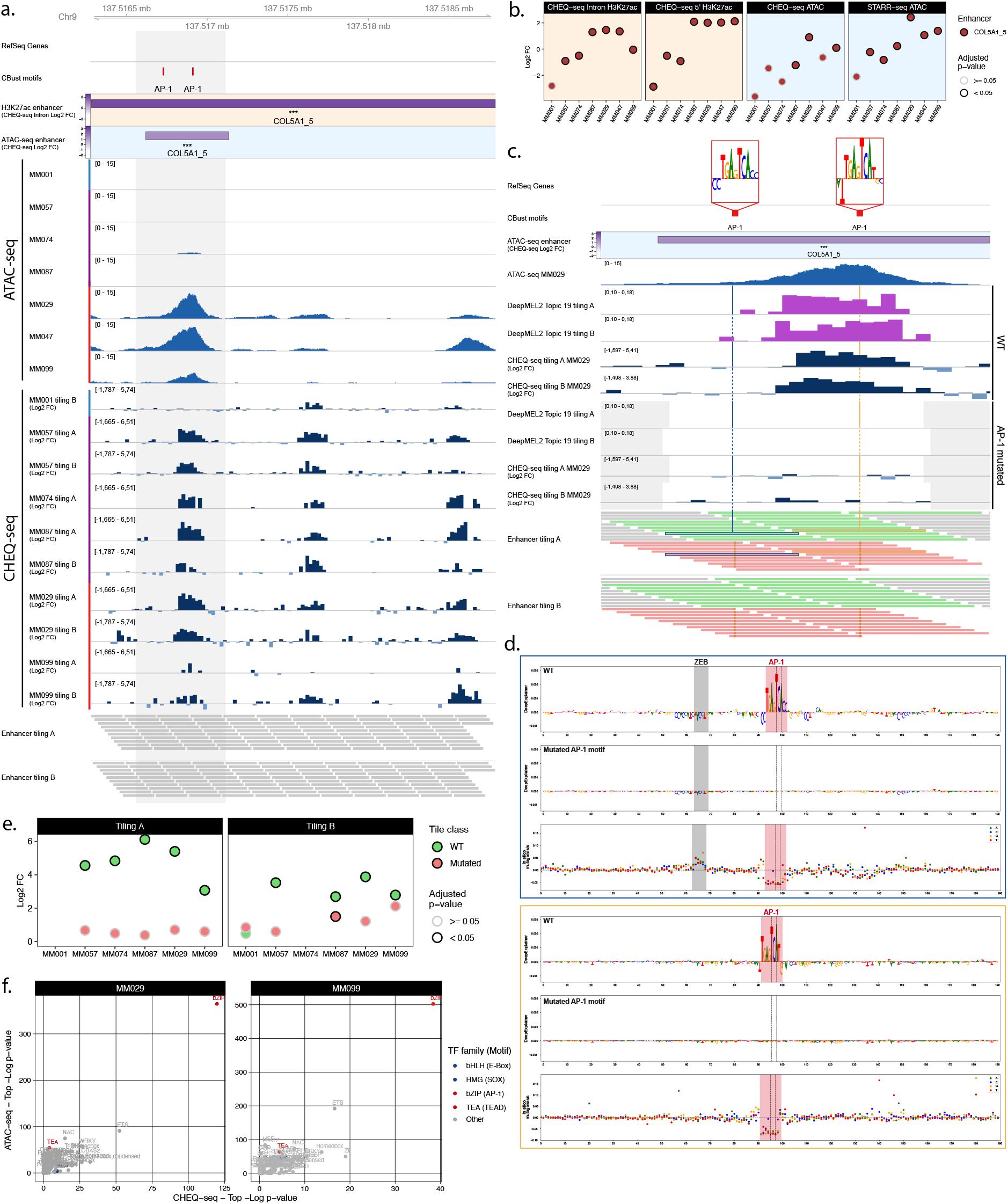
**a.**, ATAC-seq signal and CHEQ-seq WT tiles activity in the COL5A1_5 H3K27ac library region. **‘**CBust motifs’ track shows identified TFBSs for AP-1. Activity of H3K27ac and ATAC-seq regions are from MM029. **b.**, Enhancer activity for the H3K27ac ChIP-seq based and ATAC-seq based COL5A1_5 region. **c.**, CHEQ-seq activity of WT tiles and AP-1 binding site mutated tiles in MM029. ‘DeepMEL2’ tracks correspond to the accessibility prediction score for each tile. The displayed region corresponds to the highlighted area in panel **a.**. Grey areas in **‘**DeepMEL2’ and ‘CHEQ-seq tiling’ tracks correspond to regions not covered by the tiling library. In the ‘Enhancer tiling’ tracks, red and green tiles are the mutated and WT tiles, respectively. **d.**, DeepExplainer profiles for the tiles highlighted in blue and yellow in the ‘Enhancer tiling A’ track of panel **c.**. Each profile consists of the WT (top panel) and mutated (middle panel) tile nucleotide score and in silico mutagenesis score for the WT tile (bottom panel). Dashed lines indicate the location of mutated bases. **e.**, CHEQ-seq signal of the most active tile among the mutated and corresponding WT tiles for the COL5A1_5 region. **f.**, Comparison of the most enriched motif families as identified by HOMER between ATAC-seq (top ¼ most accessible tiles vs rest) and CHEQ-seq tiling libraries (all active tiles vs all inactive tiles) for MM029 and MM099.

Focusing on the first active region, enhancer activity is limited to a subset of the tiles containing an AP-1 motif (Fig 6c.). Importantly, the observed, empirical activity of a tile agrees with its DeepMEL2 prediction score. The generation of DeepExplainer profiles from those predictions reveal the presence of a ZEB repressor motif next to the first AP-1 site (Fig 6d.; Postigo and Dean, 1999). In line with this, enhancer activity is absent from all tiles containing this motif (Fig 6c.-d.). The comparison of wild type versus AP-1 mutant tiles specifically in that region further confirms the function of AP-1 as an activator in MEL intermediate and MES lines (Fig 6c.-e.). Finally, motif enrichment analysis on both the active and most accessible tiles in MES lines found only AP-1 enriched (Fig 6f.).

The gene regulatory network of the MES subtype was previously found to be mainly controlled by AP-1 and TEAD (Verfaillie et al., 2015). Our MPRA results show that only AP-1 has a direct effect on enhancer activity in the regions tested, and that the cis-regulatory role of TEAD motifs remain unknown.

## Discussion

We have investigated enhancer activity of regions specific to the two main subtypes of melanoma across a panel of patient-derived malignant melanoma lines. By studying candidate enhancers with different sequence lengths and by interpreting the results in conjunction with deep learning predictions, we obtained a better understanding of the function and specificity of melanoma enhancers.

The first studies that used MPRAs were based on very short sequences (84 to 145 nt), and identified a modular and motif-centric definition of enhancer elements (Kheradpour et al., 2013; Melnikov et al., 2012; White et al., 2013). The use of captured genomic regions allowed for longer (hundreds of bp) and more (several millions) sequences to be tested in parallel (Arnold et al., 2013; Verfaillie et al., 2016; Wang et al., 2018). While providing a genome-wide view on enhancer activity, the cloned fragments are highly variable in size, and do not allow for creating mutations. Furthermore, most studies using MPRAs have been limited to one cell line, or two very different cell lines (e.g., K562 and HepG2; or S2 and OSC in Drosophila; Arnold et al., 2013; Ernst et al., 2016), leaving the evaluation of enhancer specificity unassessed or reduced to individual examples. Here we repeat the MPRAs in a panel of seven cell lines from the same disease, allowing us to validate the cell state specificity of the selected regions.

Our approach of selecting putative enhancers based on specific H3K27ac ChIP-seq peaks, located close to differentially expressed genes, resulted in a high success rate (60 to 75%) akin to other studies leveraging a similar approach (Gorkin et al., 2020; Graybuck et al., 2021). In contrast, our large-scale library based solely on ATAC-seq peaks that decrease after SOX10 KD contained only 15.1% of active enhancers (Supplementary Fig 8). This indicates that success rates of enhancer activity vary widely, and whereas most of the differential H3K27ac peaks near marker genes are enhancers, many ATAC-seq peaks spread out across the genome, even when they are affected by TF knock-down, do not act as enhancers when tested in reporter assays. We also find active enhancers in closed chromatin regions, a phenomenon previously described in whole genome STARR-seq studies in Human and Drosophila (Arnold et al., 2013; Liu et al., 2017). Those closed regions can often be found accessible in other cell lines of our panel, and their ectopic activity suggests that the chromatin state in the genome differs from the episomal plasmid, keeping the enhancer inactive in the genome despite what seems to be a sufficient TF expression to trigger their activity in our episomal assay.

The influence of the size of the candidate enhancer has previously been assessed, although to a limited extent, and only by choosing sizes that are not representative of chromatin structures (Klein et al., 2020). Here we tested long sequences corresponding to H3K27ac regions up to 2.9 kb, 0.5 kb ATAC-seq peak regions located within the larger H3K27ac domain, and finally short 190 bp sequences tiling acetylated regions. This revealed that the actual enhancer within a H3K27ac region is usually located within the accessible sub-region of the H3K27ac region. Furthermore, the activity of that ATAC-seq peak can often be recapitulated by a sequence of 190 bp in length. Previous studies have used enhancer tiling to assess the activity of accessible chromatin regions and highlighted the importance of cell type-specific binding sites (Ernst et al., 2016; Wang et al., 2018). Nevertheless, such a short sequence does not always capture all the regulatory information found within the accessible region. Our detailed description of KIT_1A and COL5A1_5 ATAC-seq peaks highlights the presence of both activator and repressor elements spread over distances larger than 190 bp that cannot be jointly evaluated with our tiled MPRA. This finding highlights the necessity of using ∼500 bp long sequences to confidently evaluate the composition of enhancers.

Many genes are regulated by an ‘array’ of enhancers that are thought to cooperate to regulate target gene expression by forming a chromatin hub (Giammartino et al., 2020; Gorkin et al., 2014; Zhu et al., 2021). For several gene loci in this study, we selected multiple regions near the same target gene. This experiment indicated that not all regions with similar acetylation and ATAC-seq profile harbour enhancer activity in the reporter assay, but that the identification of relevant TFBSs is indicative to find active enhancers. In that regard, the use of deep learning models, even when they are trained only on ATAC-seq data, like DeepMEL2 (Atak et al., 2021), can provide a powerful tool to identify key TFs and evaluate enhancer activity.

An intriguing finding in this study is the gradient of MES enhancer activity across our panel of cell lines, with MM001, intermediate and MES lines having low, medium and high activity respectively. These enhancer profiles do not agree with their acetylation, ATAC-seq and transcriptome profiles, which all show a strong homogeneity across MEL and MES subtypes respectively. This discrepancy was more pronounced with short 190 bp sequences. Our results suggest that AP-1 is responsible for the activity of MES enhancers, agreeing with its predominance in allele-specific chromatin accessibility variants in melanoma (Atak et al., 2021). Note that AP-1 is also typically activated during stress response (Hess et al., 2004). This may explain the limited activity of MES regions in MM001 following electroporation. Nevertheless, AP-1 enhancers show much higher activity observed in intermediate lines, which cannot be explained by a stress response only. The observed activity of AP-1 enhancers in intermediate lines, even though they are not accessible, nor acetylated in these lines, could be due to the episomal reporter assay, whereby the (limited) AP-1 activity in these lines can activate enhancers on a plasmid, but not in the endogenous locus. The use of a genome integrated reporter assay could help to determine if the accessibility of the sequence is required for MES enhancer activity in MEL lines.

The analysis of MEL regions revealed that both SOX and MITF binding sites independently contribute to enhancer activity. TFAP2 binding sites did not show any influence on enhancer activity in the tested H3K27ac regions and showed only a small enrichment (NES: 1.19, pAdj: 0.046) in SOX10 dependent ATAC-seq peaks displaying activity. Those results contradict previous observations of TFAP2 and MITF co-occurrence in active regulatory elements in melanocytes and of TFAP2 pioneer function during cell fate commitment in neural crest cells (Rothstein and Simoes-Costa, 2020; Seberg et al., 2017). These differences may be explained at least in part by the limited number of regions selected in our H3K27ac library or by a possible reduced influence of TFAP2 in the MM lines used in this study. Interestingly, we identify an interaction between SOX and MITF where the presence of at least one SOX dimer motif greatly increases the activity of enhancers containing MITF motifs. It is not clear if this interaction is due to the direct cooperation of SOX10 and MITF or the known pioneer factor function of SOX family members (Julian et al., 2017). The cooperation between these two TFs could be governed by clear rules requiring the presence of a SOX motif for activity or by the necessity of interactions through heterotypic clusters of TFBSs to maintain a high level of activity. Indeed, studies have shown that, in some cases, homotypic clusters of TFBSs for a single TF can drive lower expression than heterotypic clusters of TFBSs for multiple TFs (Fiore and Cohen, 2016; Levo and Segal, 2014; White et al., 2016). Our synthetic combinations assay did not include sequences with less than 6 MITF motifs without SOX motifs but we did identify such type of sequences active in our enhancer tiling assay (e.g. KIT_1 and SOX10_4 Supplementary Fig 3b.-c.), suggesting that a SOX motif is not required in enhancers with few MITF motifs. Alternatively, the pioneer function of SOX would suggest that the sequences cloned in the reporter vector are subject to, at least partial, chromatinisation. Riu et al. previously presented evidence supporting plasmid chromatinisation, more precisely of regions from bacterial origin resulting in silencing of a human expression cassette (Riu et al., 2007). However, it is still unclear if non-coding human regions could also be subject to chromatinisation in a plasmid context and what role sequence length might play. We find a less pronounced influence of the SOX motif in short 190 bp sequences compared to 259 bp sequences, and shorter MES enhancers also show more ectopic activation in intermediate lines, which may suggest that short enhancer sequences are being less efficiently chromatinised. Consequently, short enhancers might be characterised by increased accessibility in an extrachromosomal reporter, uncoupling measured activities from those demonstrated at the endogenous locus. Kheradpour et al. proposed a similar hypothesis by suggesting that DNA sequence features contained within the tested elements are partly responsible for establishing the endogenous chromatin state (Kheradpour et al., 2013). These considerations are also relevant in the context of extrachromosomal DNA driving oncogene overexpression in various human malignancies (Verhaak et al., 2019). The use of integrated MPRA or the measurement of chromatin accessibility in the plasmid would help to determine if enhancer sequences are subject to chromatin modification.

With this study we show that melanoma subtype specific enhancers can be identified, even to the size of 190bp, which indicates promising avenues to use such information gathered from MPRAs to identify small and specific enhancers for enhancer therapy. Our assay with combinations of SOX and MITF motifs showed that new MEL enhancers can be designed by addition of TFBSs to a random background sequence and that the level of activity can be controlled by varying the number of TFBSs. Using genomic regions confirmed as MEL enhancers and adjusting the motif sequences and numbers could further improve enhancer design.

## Methods

### MPRA design and cloning

#### H3K27ac based library

The selection of H3K27ac ChIP-seq regions harbouring potential state-specific enhancer activity was done as follows. Differentially enriched H3K27ac ChIP-seq peaks between MEL and MES state, identified in our previous study (Verfaillie et al., 2015), were used as input in i-cisTarget (Imrichová et al., 2015) to identify enriched TFBSs in specifically acetylated regions. For each melanoma subtype, the target genes of the top 3 enriched TFBSs were extracted and used to filter the list of differentially expressed genes identified in each subtype (Verfaillie et al., 2015). From the remaining top differentially expressed genes, ChIP-seq tracks are visually investigated using the UCSC Genome Browser for regions displaying acetylation peaks only in one state close to those genes (Regions of ∼100 kb upstream and downstream of the genes were explored). Candidate regions were preferentially selected if they also overlapped with ChIP-seq peaks for SOX10 or MITF for MEL regions and JUN and JUNB (AP-1) for MES regions. 65 regions with a size < 3 kb were manually selected and primers to amplify them were designed using Primer3Plus within the flanking first 100 bp on each side of the sequence. 15 bp extensions were added to the primers to allow recombination with the region upstream of the SCP1 promoter in the CHEQ-seq vector (Verfaillie et al., 2016) via In-Fusion reaction (Primer list in Supplementary Table 2).

All PCR amplifications performed in this study make use of the KAPA HiFi HotStart ReadyMix (Roche, Basel, Switzerland). The amplification of the selected regions was done from MM074 genomic DNA in a 50 µl reaction. PCR fragments were then purified on a 0.8% agarose gel and the sequence was confirmed via Sanger sequencing. 53 regions could be successfully amplified (Supplementary Table 1). For the generation of the CHEQ-seq 5’ library, the CHEQ-seq vector containing a random 17 bp barcode (BC) upstream of the synthetic intron was linearized by inverse PCR with the primers “CHEQ-seq_lin_5’_For” and “CHEQ-seq_lin_5’_Rev” resulting in a fragment with both ends overlapping with the primers designed to amplify the selected regions. Amplified regions were mixed in equimolar ratio and introduced in the CHEQseq vector via In-Fusion reaction (Takara Bio, Kusatsu, Japan) with a vector to insert ratio of 1:2.

For the generation of the CHEQ-seq Intron library, amplified regions were reamplified with primers containing adaptors to allow recombination within the intron of the CHEQ-seq vector (“H3K27Ac_lib_intron_For”, “H3K27Ac_lib_intron_Rev”). The CHEQ-seq vector containing a random 18 bp BC downstream of the synthetic intron was linearized by inverse PCR with the primers “CHEQ-seq_lin_Intron_For” and “CHEQ-seq_lin_Intron_Rev”. Reamplified regions were mixed in equimolar ratio and introduced in the CHEQseq vector via In-Fusion reaction with a vector to insert ratio of 1:2. The In-Fusion reactions were dialysed against water in a 6 cm petri-dish with a membrane filter MF-Millipore 0.05 µm (Merck, Kenilworth, New Jersey) for 1 hour. Reactions are recovered from the membrane and 2.5 µl of the reaction are transformed into 25 µl of Lucigen Endura ElectroCompetent Cells (Biosearch Technologies, Hoddesdon, United Kingdom). Transformed bacteria are cultured overnight in a shaker before maxiprep.

### ATAC-seq based library

The sequences constituting the ATAC-seq based library are selected from ATAC-seq peaks from MM001, MM029, MM047, MM057, MM074, MM087 and MM099 overlapping with H3K27ac library regions. Among this subset, only peaks that overlap with regions identified as active in the H3K27ac library or that are assigned as MEL-or MES-specific regulatory regions (respectively represented by regions belonging to topic 4 and topic 7) in our previously published cisTopic analysis of ATAC-seq data from 16 human melanoma cell lines were retained (Bravo González-Blas et al., 2019). The final selection included 46 ATAC-seq peaks (Supplementary Table 1). The summit of each peak was extended by 250 bp on both sides to generate a 501 bp sequence. At the 3’ end of the sequence a 16 bp spacer sequence was added, followed by an 8 bp BC specific for each selected region. 38 sequences were synthesised by TWIST Biosciences (South San Francisco, California). This includes 4 sequences where an A or a T was substituted to break a polyA or polyT respectively in order to make the sequences compatible with the synthesis process. Primers were designed for the remaining 8 sequences to amplify them from genomic DNA (“ATAC based library amp” in Supplementary table x). Only one PCR, i.e. for the amplification of the SERPINE1_2 sequence, was not successful. The CHEQ-seq vector containing a random 17 bp BC upstream of the synthetic intron and the STARR-seq ORI vector (addgene #99296; Muerdter et al., 2017) were linearised via inverse PCR (“ATAC based library cloning” in Supplementary table 2). The individual sequences were pooled together in equimolar ratio and NEBuilder (New England Biolabs, Ipswich, Massachusetts), with a vector to insert ratio of 1:2, was used to introduce them in the vectors. Dialysis and transformation are performed similarly to the H3K27ac library. Before culture for maxiprep, 1:100,000 of the transformed bacteria is plated on a LB-agar dish with carbenicillin to estimate the complexity of the cloned library. A volume of bacteria corresponding to a complexity of 3,500 BCs per enhancer is put in culture for maxiprep.

#### Enhancer tiling libraries

Using a custom script, tiles were generated by selecting 190 bp from the start (library A) or position 11 (library B) of the H3K27ac selected regions and by switching every 20 bp in 3’ direction. Tile generation is stopped when the position of the final nucleotide of the tile is superior to the final nucleotide of the H3K27ac region. To generate mutated tiles, we first scanned all selected H3K27ac regions with Cluster-Buster (cbust; Frith et al., 2003) and the following position weight matrices (PWM) separately: transfac_pro M08838 (SOX10 dimer); homer RTCATGTGAC_MITF; transfac_pro M01859 (TFAP monomer); tfdimers MD00038 (TFAP dimer); tfdimers MD00591 (SOX10-TFAP dimer); hocomoco JUN_f1; homer NATGASTCABNN_Fosl2; cisbp M5907 (TEAD). Using a custom script, motifs were mutated on their 2 or 4 most important nucleotides for monomers and dimers respectively. Mutated tiles for each motif were generated separately so that all occurrences of a single motif are present. Shuffled negative control tiles were generated by shuffling all wild type and mutated sequences with ushuffle (Jiang et al., 2008). Sequences containing a stretch of the same nucleotide for 6 or more nucleotides were filtered out. The remaining tiles were scored with cbust using the same PWMs and parameters as before and 800 tiles containing no motifs were selected at random. In total, libraries A and B contained 7412 and 7393 tiles respectively (Supplementary Table 1).

Adaptor sequences were added to the tiles: “Adaptor_LibA_5’” and “Adaptor_LibA_3’” for library A and “Adaptor_LibB_5’” and “Adaptor_LibB_3’” for library B. The use of different adaptors for each library will result in the insertion of the library B 20 bp downstream of the library A, providing a slightly different surrounding context which combined with a high number of barcodes per enhancer aim at reducing experimental noise. Final libraries were synthesised via Agilent’s Oligonucleotide Library Synthesis Technology (Santa Clara, California).

Oligonucleotides libraries were resuspended in endotoxin free TE buffer pH 8 to a final concentration of 20 nM. For each library, 10 PCR reactions are performed with 2 µl of resuspended librarie for 12 cycles with primers “Lib_A_amp_For” and “Lib_A_amp_Rev” or “Lib_B_amp_For” and“Lib_B_amp_Rev”. The PCR product was first cleaned up using MinElute (Qiagen) with 5 PCR reactions per column then pooled together and cleaned up a second time with 1.6X SPRI beads (Beckman Coulter, Brea, California). The CHEQ-seq vector containing a random 17 bp BC upstream of the synthetic intron was linearised via inverse PCR with primers “CHEQseq_lin_A_For” and “CHEQseq_lin_A_Rev” or “CHEQseq_lin_B_For” and “CHEQseq_lin_B_Rev”. Amplified libraries and the corresponding linearised vector were combined in an NEBuilder reaction with a vector to insert ratio of 1:3.25. Dialysis, transformation and maxiprep are performed similarly to the H3K27ac library.

#### SOX10 knockdown based library

To define a set of SOX10-dependent MEL enhancers, we used public OmniATAC-seq data during a time series of SOX10 knockdown (0, 24, 48, 72h) on two melanoma cells lines (MM057 and MM087) (GSE114557; Bravo González-Blas et al., 2019). We generated 50 simulated single cells per condition by randomly sampling 50,000 reads per cell. Candidate regulator regions were defined by peak calling with MACS2 (v.2.0.10) in each of the bulk samples and by merging the condition-specific peaks using mergeBed (part of BEDtools, v.2.23.0), and blacklisted regions were removed using https://sites.google.com/site/anshulkundaje/projects/blacklists (hg19). We ran cisTopic (v0.2.1; Bravo González-Blas et al., 2019) (parameters: α = 50/T, β = 0.1, burn-in iterations = 500, recording iterations = 1,000) for models with a number of topics between 2 and 25. The best model was selected on the basis of the highest log-likelihood, resulting in 16 topics. We binarized the topics using a probability threshold of 0.99 and continued with topic 11, which contained regions that are accessible in baseline conditions but lose accessibility following SOX10 knockdown. A total of 1,461 enhancers were chosen to be tested. To increase the resolution of the library, we tiled the sequences to 190 bp using a 120 bp sliding window across the enhancers resulting in 6696 tiles. In addition, 100 shuffled negative control sequences were generated similarly to the tiling libraries (Supplementary Table 1). Vector specific adapters were added to the sequences and the tiles were synthesised via Agilent’s Oligonucleotide Library Synthesis Technology.

The library was amplified via PCR using primers “Lib_A_amp_For” and “Lib_A_amp_Rev” and cloned into the linearized CHEQ-seq plasmid by following the same procedure as for the enhancer tiling library A. Dialysis, transformation and maxiprep are performed similarly to the H3K27ac library.

#### Synthetic combinations of SOX and MITF library

Random 259 bp sequences were generated using SMS2 - Random DNA Sequence tool (Stothard, 2000). Two sequences, displaying no enrichment in any of the topics defined in our previously published cisTopic analysis (Bravo González-Blas et al., 2019), were selected as background sequences. SOX and MITF motifs (ACAAAGACGGCTTTGT and CACGTG respectively) are inserted in the sequence with a motif in the centre and the other motifs placed upstream and downstream with a distance of 25, 50 or 75 bp (Fig 5e. top panel). A complete list of motif combinations can be found in Supplementary table 1. Two hundred negative control shuffled sequences are generated with ushuffle as described previously. An 11 bp BC specific for each enhancer is placed in 5’ position of the sequence. Barcoded enhancers are finally flanked with the adaptors GAGCATGCACCGGTG and CGCTTCGAGCAGACA in 5’ and 3’ respectively. The final library was synthesised by Twist Bioscience as an Oligo Pool. The oligonucleotide library is resuspended according to manufacturer recommendation and amplified via PCR with the primers “CHEQ_comb_amp_For” and “CHEQ_comb_amp_Rev”. The CHEQ-seq vector with a 17 bp random BC upstream of the intron is linearised via inverted PCR with the primers “CHEQ_comb_lin_For” and “CHEQ_comb_lin_Rev”. NEBuilder is then used to combine the library with the linearised vector with a vector to insert ratio of 1:2. Dialysis and transformation are performed similarly to the H3K27ac library. Before culture for maxiprep, 1:100,000 of the transformed bacteria is plated on a LB-agar dish with carbenicillin to estimate the complexity of the cloned library. A volume of bacteria corresponding to a complexity of 500 BCs per enhancer is put in culture for maxiprep.

### Enhancer – Barcode assignment

#### CHEQ-seq 5’/Intron for H3K27ac ChIP-seq regions

The part of the plasmid extending from the enhancer till the random BC is amplified via PCR with primers “PacBio_5’_For”, “PacBio_5’_Rev”, “PacBio_Intron_For” and “PacBio_Intron_Rev” for CHEQ-seq 5’ and CHEQ-seq Intron respectively. Gel extraction is performed to isolate the PCR product with the correct size range using the NucleoSpin Gel and PCR Clean-up kit (Macherey-Nagel, Düren, Germany). PacBio sequencing library preparation for both libraries was done by the Genomics Core Leuven (KU Leuven). Sequencing was done with a PacBio Sequel for long-read sequencing (Pacific Biosciences, Menlo Park, California) with both libraries sequenced with one SMRT cell. We obtained 66,407 and 105,590 reads with 3 passes for CHEQ-seq 5’ and CHEQ-seq Intron respectively.

Enhancer - BC assignment was done with a custom script. Briefly, enhancers and random BCs were independently extracted from the reads with Cutadapt (Martin, 2011). Enhancer sequences are mapped with Minimap2 (Li, 2018) and a custom genome containing all the cloned regions, only MAPQ>=4 were retained. Mapped enhancers were linked back to random BCs of the vector. Following assignment, 46 (86.8%) and 50 (94.3%) sequences could be identified in the CHEQ-seq 5’ and Intron library respectively. The CHEQseq 5’ library displayed an average of 31.9 BCs per enhancer while the Intron library displayed an average of 604.5 BCs per enhancer.

#### CHEQ-seq for ATAC-seq regions

A PCR amplification of the enhancer specific BC together with the random BC is done with the primers “Enh-BC_ATAC_Stag=X_For” and “Enh-BC_ATAC_Rev”. Illumina sequencing adaptors are added during a second round of PCR with the primers “i5_Indexing_For” and “i7_Indexing_Rev”. After sequencing in NovaSeq600 for 50 cycles in read 1 and 49 cycles in read 2, enhancer BCs and random BCs are extracted from read 1 and read 2 respectively with Cutadapt before being filtered for quality (Q>30). Following assignment, 44 (95.7%) sequences could be identified, with an average of 34,815 BCs per enhancer.

#### CHEQ-seq for enhancer tiling/SOX10-KD library

Cloned sub-libraries are amplified via PCR with the primers “Enh-BC_Tiling-A_Stag=X_For” (for sub-library A and SOX10-KD library), “Enh-BC_Tiling-B_Stag=X_For” (for sub-library B) and “Enh-BC_Tiling_Rev”. Illumina sequencing adaptors are added during a second round of PCR with the primers “i5_Indexing_For” and “i7_Indexing_Rev”. After sequencing in NovaSeq600 for 251 cycles in read 1 and 51 cycles in read 2, whole length enhancers and random BCs are extracted from read 1 and read 2 respectively with Cutadapt before being filtered for quality (Q>30). Enhancer reads are mapped and linked to random BCs as previously described for CHEQ-seq 5’/Intron for H3K27ac ChIP-seq regions. Following assignment, 7,356 (99.2%), 7,344 (99.8%) and 6,773 (99.7%) sequences could be identified in the sub-libraries A and B and the SOX10-KD library respectively, with an average of 3,096, 3,021 and 8,056 BCs per enhancer.

#### CHEQ-seq for SOX-MITF synthetic combinations

This CHEQ-seq library is prepared for sequencing similarly to the CHEQ-seq enhancer tiling A sub-library. After sequencing in NovaSeq600 for 50 cycles in read 1 and 49 cycles in read 2, enhancer BCs and random BCs are extracted from read 1 and read 2 respectively with Cutadapt before being filtered for quality (Q>30). Following assignment, 249 (99.6%) sequences could be identified, with an average of 276 BCs per enhancer.

### Cell culture

All MM lines were cultured in Ham’s F10 nutrient mix (ThermoFisher Scientific, Waltham, Massachusetts) supplemented with 10% fetal bovine serum (ThermoFisher Scientific) and 50 μg ml-1 penicillin/streptomycin (ThermoFisher Scientific). Cell cultures were kept at 37°C, with 5% CO_2_.

### MPRA assay

The MPRA libraries were electroporated in 4 to 6 million cells each using the Nucleofector 2b or 4D (Lonza, Basel, Switzerland) with 6 µg of plasmid DNA and program T-030 or Y-001 and DS-132, EH-116 or CM-134 respectively. For the CHEQ-seq 5’ and Intron H3K27ac and CHEQ-seq ATAC libraries, one replicate was performed per cell line except for MM087 where three replicates were performed. For STARR-seq ATAC and the CHEQ-seq SOX-MITF combinations library, one replicate was performed per cell line. For the CHEQ-seq enhancer tiling libraries, two replicates were performed per cell line except for MM087 where four replicates were performed. For the CHEQ-seq SOX10-KD library, three replicates were performed in MM087. Medium was changed 24 hours after electroporation. 48 hours post-electroporation, cells were detached from the plate using trypsin (ThermoFisher Scientific). One fifth of the cells was used for plasmid DNA extraction (Qiagen, Hilden, Germany). The remaining cells underwent RNA extraction using the innuPREP RNA Mini Kit 2.0 (Analytik Jena, Jena, Germany), followed by mRNA isolation using the Dynabeads mRNA purification kit (Ambion, Austin, Texas) and cDNA synthesis using the GoScript RT kit and oligo dT primer (Promega, Madison, Wisconsin). For STARR-seq samples, a junction PCR is done for 12 cycles with the primers “STARR-seq_Junction_cDNA_For” or “STARR-seq_Junction_plasmid_For” and “STARR-seq_Junction_Rev” followed by a PCR to amplify the enhancer for 4 cycles with the primers “STARR-seq_enhancer_Stag=X_For” and “STARR-seq_enhancer_Rev”. For CHEQ-seq samples, a PCR was performed to amplify the random BC from the plasmid DNA or cDNA samples for 16 cycles with the primers pairs “Cheq-seq_barcode_Intron_round1_Stag=0_For”/”Cheq-seq_barcode_Intron_round1_Rev” for the CHEQ-seq Intron H3K27ac library or “Cheq-seq_barcode_5’_round1_Stag=X_For”/”Cheq-seq_barcode_5’_round1_Rev” for all other libraries. To add Illumina sequencing adaptors, all samples were finally amplified by PCR for 6 cycles with the primers “i5_Indexing_For” and “i7_Indexing_Rev”. After confirmation of the fragment size with a Bioanalyzer, samples were sequenced at the Genomics Core Leuven (KU Leuven).

### MPRA analysis

#### Read processing and BC identification

Read processing following sequencing is performed with a custom bash script. First, random BCs are extracted using Cutadapt and filtered for a quality (Q-score) > 30. The number of reads per uniquely identified BC is counted and the name of the enhancer is assigned to the BC sequence based on the enhancer - BC assignment list for each library. Unassigned BCs are filtered out to obtain a final data frame containing the name of the enhancer, the BC sequence and the number of reads.

#### Estimation of enhancer activity

Enhancer activity from MPRA assay is estimated via a custom R script (RStudio, R version 3.6.0). Enhancers are first filtered based on the number of BCs identified in the sequencing reads. Thresholds of 5 (for the H3K37ac libraries and enhancer tiling libraries), 10 (for ATAC libraries) or 20 (for the SOX-MITF combinations library) BCs per enhancer are selected based on the complexity of the library and the sequencing saturation of the enhancer - BC assignment samples. For the remaining enhancers, BC counts are aggregated per enhancer and then a count per million (CPM) normalisation is applied. Plasmid (input) and cDNA (output) samples are merged by keeping only enhancers remaining in both samples after filtering. Input normalisation is done by dividing CPM normalised cDNA counts by CPM normalised plasmid counts resulting in a FC value. For libraries with shuffled sequences, a basal expression normalisation is further applied by dividing the FC value of the enhancer by the median FC value of the shuffled sequences. The MPRAnalyze method was tried as an alternative to our method and gave nearly identical results in the case of the H3K27ac CHEQ-seq Intron library (mean Pearson’s correlation on log2 FC r = 0.96; Ashuach et al., 2019). The high computational demand when the number of BCs is high made it inappropriate for the analysis of most libraries. For consistency, all assays were analysed with our aggregated BC method which showed more consistency with low complexity libraries and more scalability with very complex libraries. In order to distinguish active and inactive enhancers, a Gaussian fit of the shuffled negative control values is performed with the “robustbase 0.93-6” package and a p-value and Benjamini–Hochberg adjusted p-value is calculated based on that Gaussian fit for all enhancers with the “stats 3.6.0” package. An enhancer is considered active if its adjusted p-value is < 0.05. For the H3K27ac and ATAC-seq libraries, that did not contain shuffled sequences, regions containing no active tiles in the enhancer tiling MPRAs and displaying low activity in both H3K27ac and ATAC-seq libraries MPRAs are selected as negative controls and used to fit the Gaussian curve. For the CHEQ-seq SOX10-KD library, DEseq2 (Love et al., 2014) was used for estimating enhancer activity.

#### Sample exclusion

Despite the high number of identified enhancer - random barcode couples during the assignment step for the CHEQ-seq enhancer tiling libraries A and B, the complexity of those libraries was so high that less than 3% of the barcodes could be identified following MPRA assay. This resulted in a low enhancer coverage or an insufficient number of remaining reads to identify enhancer activity in many samples. The “OutlierD 1.48.0” R package was used to identify outliers. Samples which displayed <1% of outliers or had too low coverage (<450 tiles) were excluded.

### Motif enrichment and GSEA analysis

Differential motif enrichment between the active tiles and the remaining tiles for the enhancer tiling libraries and the SOX10-KD library was performed via Homer findMotifs (Heinz et al., 2010). For the enhancer tiling libraries, differential motif enrichment was also performed between the top 1/4th accessible tiles and the remaining tiles. MITF, SOX10 and TFAP2A ChIP-seq peaks (Laurette et al., 2015; Seberg et al., 2017) were intersected with the tiles using BEDtools. A GSEA analysis was performed using the R package “fgsea” (Korotkevich et al., 2021) by ranking the tiles according to their log2 FC and providing the overlaps of the ChIP-seq peaks with the tiles as gene sets.

### Deep learning predictions and nucleotide contribution visualisation

Enhancer sequences for the enhancer tiling and SOX-MITF combinations libraries are scored with the DeepMEL2+GABPA version of DeepMEL2 as described previously (Atak et al., 2021). To accommodate for the 500 bp required length of the sequences to be scored by DeepMEL2, the vector sequence flanking the insertion site of the enhancer is added on both sides of the sequence. A threshold of 0.1 was defined to distinguish between low and high prediction score for topics 16, 17 and 19 as it approaches the mean score + 2 * standard deviation of those topics.

Visualisation of nucleotide contribution to DeepMEL2 prediction score is done with DeepExplainer as described previously (Atak et al., 2021; Lundberg and Lee, 2017).

### ChIP-seq, ATAC-seq and RNA-seq public data

MITF and SOX10 ChIP-seq in 501mel were downloaded from the GEO entry GSE61965 (Laurette et al., 2015). TFAP2A ChIP-seq in human primary melanocytes purified from discarded neonatal foreskin samples was downloaded from the GEO entry GSE67555 (Seberg et al., 2017). JUN and JUNB ChIP-seq in MM099 line were downloaded from the GEO entry GSE159965 (Atak et al., 2021). H3K27ac ChIP-seq for MM001, MM011, MM031, MM034, MM047, MM057, MM074, MM087, MM099, MM118, SKMEL5 were downloaded from the GEO entries GSE60666 and GSE114557 (Bravo González-Blas et al., 2019; Verfaillie et al., 2015). OmniATAC-seq data for MM001, MM011, MM029, MM031, MM034, MM047, MM057, MM074, MM087, MM099 and MM118 were downloaded from the GEO entries GSE142238 and GSE134432 (Minnoye et al., 2020; Wouters et al., 2020). SOX10-KD time course OmniATAC-seq for MM057 and MM087 were downloaded from the GEO entry GSE114557 (Bravo González-Blas et al., 2019). Single cell RNA-seq data for MM001, MM029, MM047, MM057, MM074, MM087 and MM099 were downloaded from the GEO entry GSE134432 (Wouters et al., 2020).

### Data access

MPRA data generated for this study have been submitted to the NCBI Gene Expression Omnibus (GEO, https://www.ncbi.nlm.nih.gov/geo/) under accession number GSE180879.

## Supporting information

Supplementary Table 1 - Libraries composition

Supplementary Table 2 - Primer list

## Acknowledgements

This work was supported by an ERC Consolidator Grant to S.A. (no. 724226_cis-CONTROL), by the KU Leuven (grant no. C14/18/092 to S.A.), by the Foundation Against Cancer (grant no, 2016-070 to S.A.), a PhD and a postdoctoral fellowship from the FWO (L.M., no. 1S03317N, J.D. no. 12J6916N, respectively) and a postdoctoral research fellowship from Kom op tegen Kanker (Stand up to Cancer), the Flemish Cancer Society, and from Stichting tegen Kanker (Foundation against Cancer), the Belgian Cancer Society (J.W.). Computing was performed at the Vlaams Supercomputer Center and high-throughput sequencing via the Genomics Core Leuven. MM lines were a kind gift from Pr. Ghanem-Elias Ghanem (Institut Jules Bordet, ULB, Belgium). The funders had no role in study design, data collection and analysis, decision to publish or preparation of the manuscript.

## Supplementary figures

**Supplementary figure 1:**
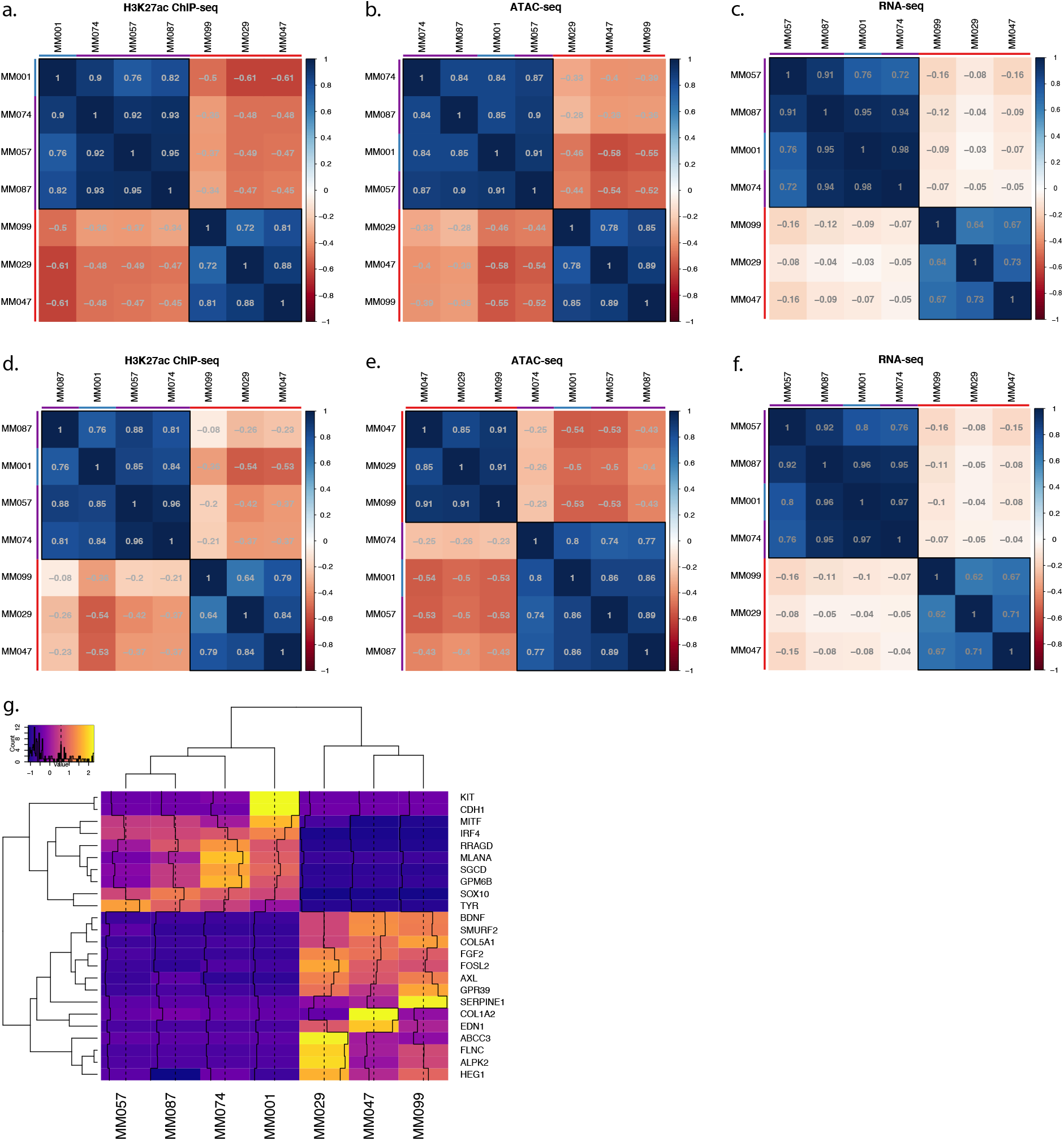
**a.-c.**, Correlation tables displaying the Pearson correlation coefficient for H3K27ac ChIP-seq mean signal of the H3K27ac library regions (**a.**), ATAC-seq mean signal of the same regions (**b.**) and gene expression of the predicted region’s target genes from single-cell RNA-seq (**c.**). **d.-f.**, Same correlation tables as **a.-c**. but for the ATAC-seq based library regions. Red, purple and blue bars indicate MES, Intermediate and MEL lines respectively. **g.**, Mean transcript expression from single-cell RNA-seq data of each gene associated with an enhancer.

**Supplementary figure 2:**
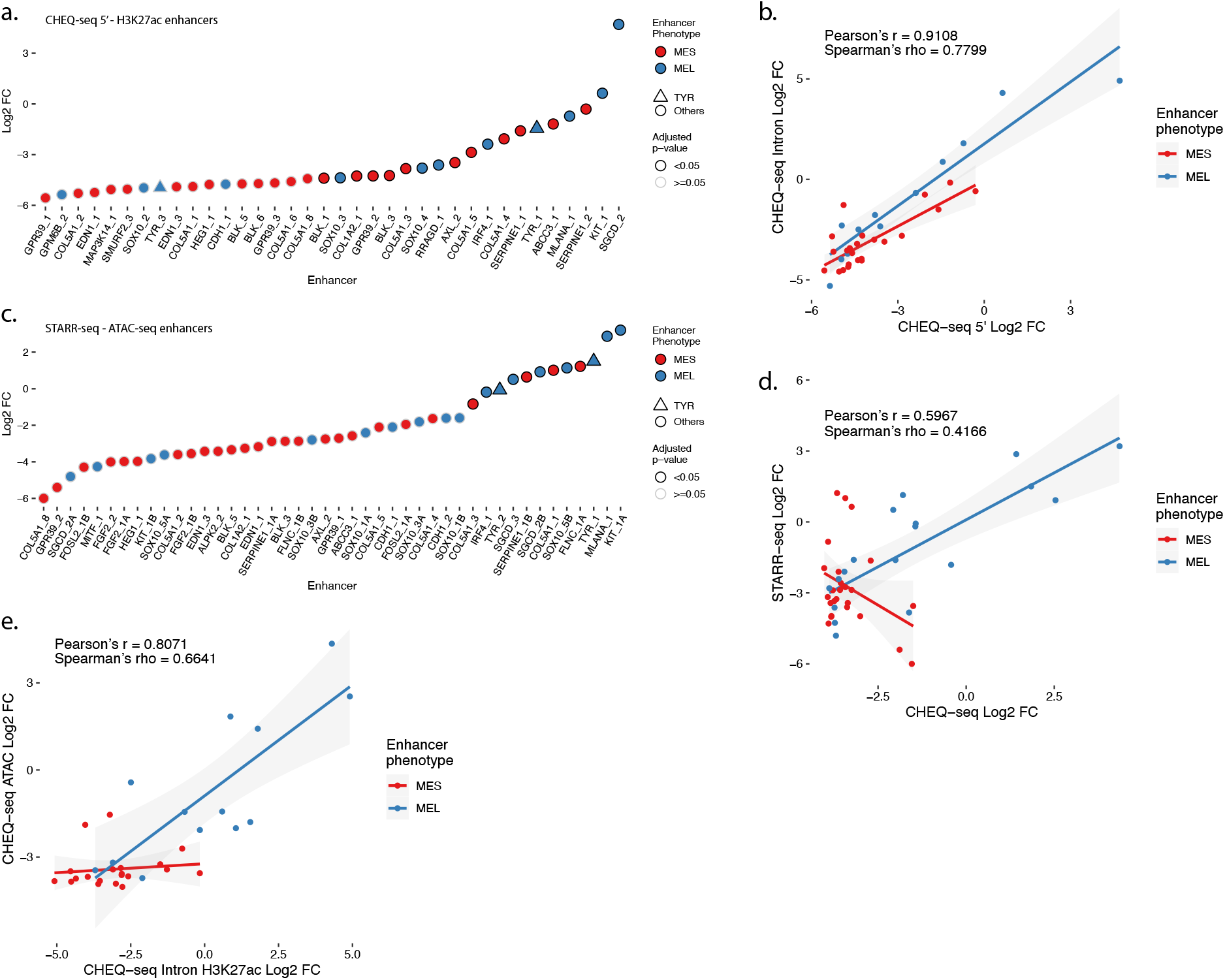
**a.**, Enhancer activity profile for the CHEQ-seq intron H3K27ac library in MM001. **b.**, Scatterplot representation of CHEQ-seq intron vs CHEQ-seq 5’ H3K27ac library in MM001. **c.**, Enhancer activity profile for the STARR-seq ATAC-seq library in MM001. **d.**, Scatterplot representation of CHEQ-seq vs STARR-seq ATAC-seq library in MM001. **e.**, Scatterplot representation of CHEQ-seq intron H3K27ac library vs CHEQ-seq ATAC-seq library in MM001. For H3K27ac regions with 2 ATAC-seq peaks, the highest value was assigned to the region.

**Supplementary figure 3:**
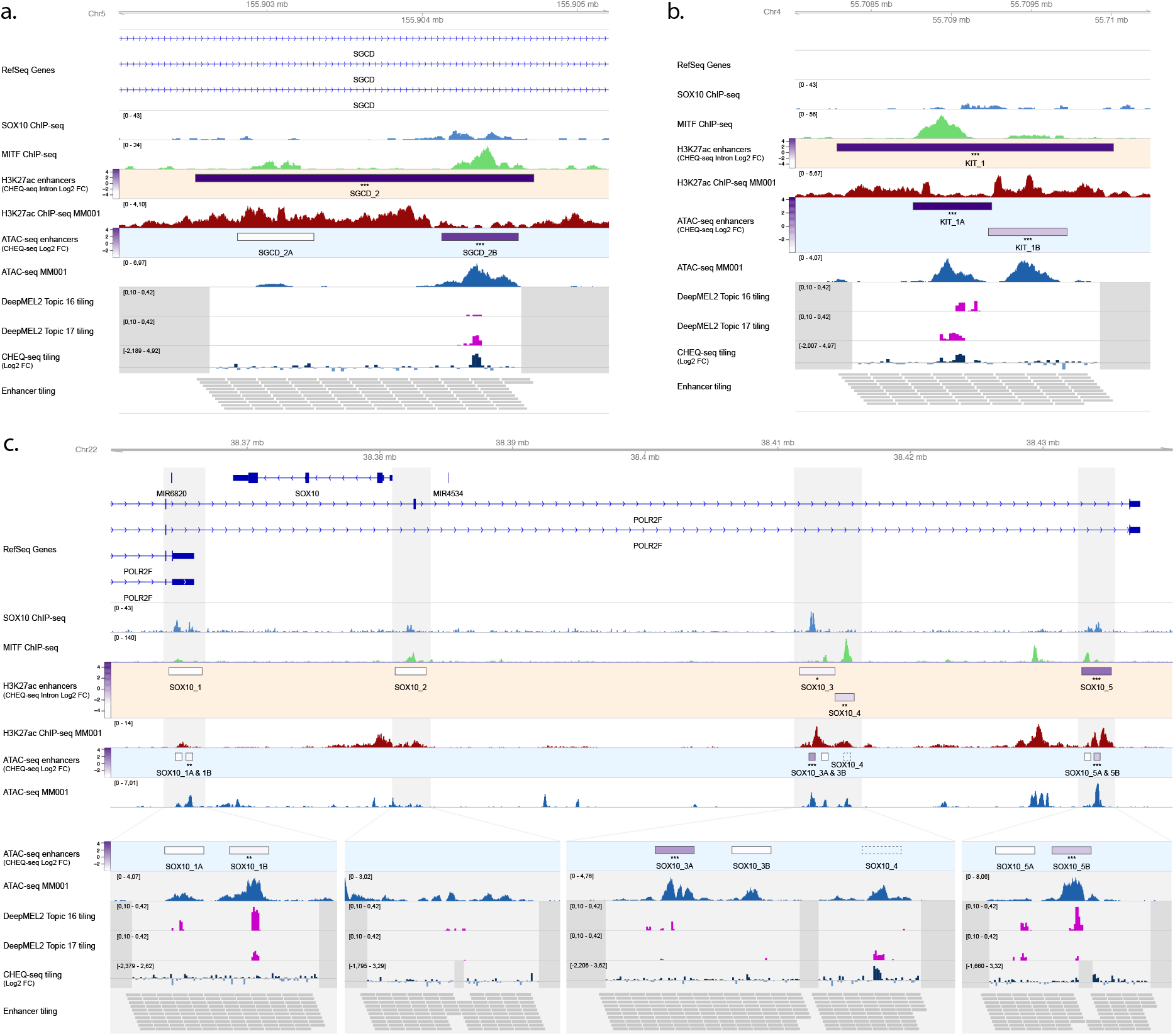
**a.**, **b.**, Enhancer activity of SGCD_2 (**a.**), KIT_1 (**b.**) and SOX10 (**c.**) regions. SOX10 and MITF ChIP-seq, H3K27ac ChIP-seq and ATAC-seq for MM001 and DeepMEL2 predictions and CHEQ-seq values of the enhancer tiling are displayed. Dark grey areas are regions not covered by the tiling library. CHEQ-seq activity is visible in the **H3K27ac enhancers** and **ATAC-seq enhancers** tracks. Benjamini–Hochberg adjusted p-values: * < 0.05; ** < 0.005; *** < 0.001.

**Supplementary figure 4:**
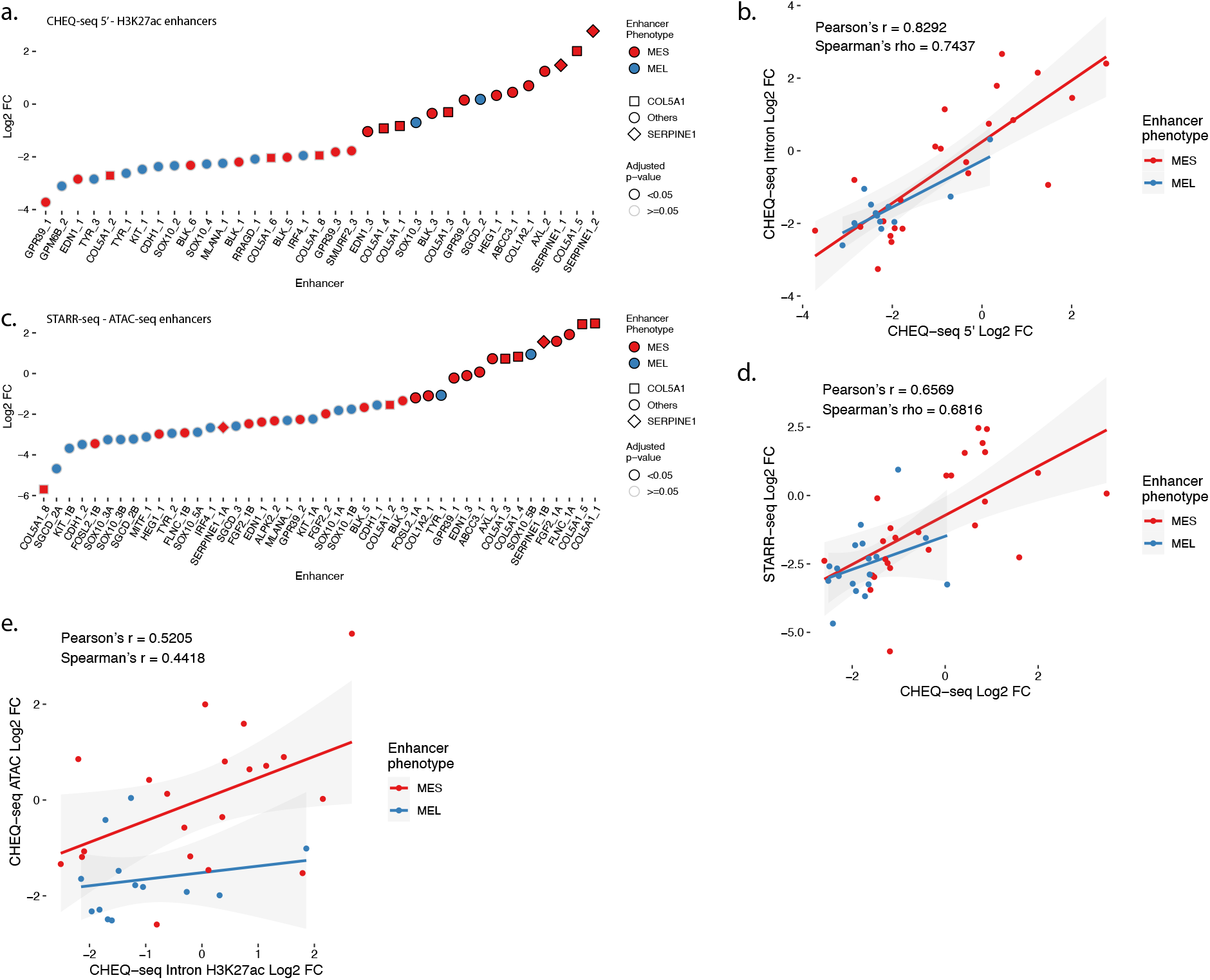
**a.**, Enhancer activity profile for the CHEQ-seq intron H3K27ac library in MM029. **b.**, Scatterplot representation of CHEQ-seq intron vs CHEQ-seq 5’ H3K27ac library in MM029. **c.**, Enhancer activity profile for the STARR-seq ATAC-seq library in MM029. **d.**, Scatterplot representation of CHEQ-seq vs STARR-seq ATAC-seq library in MM029. **e.**, Scatterplot representation of CHEQ-seq intron H3K27ac library vs CHEQ-seq ATAC-seq library in MM029. For H3K27ac regions with 2 ATAC-seq peaks, the highest value was assigned to the region.

**Supplementary figure 5:**
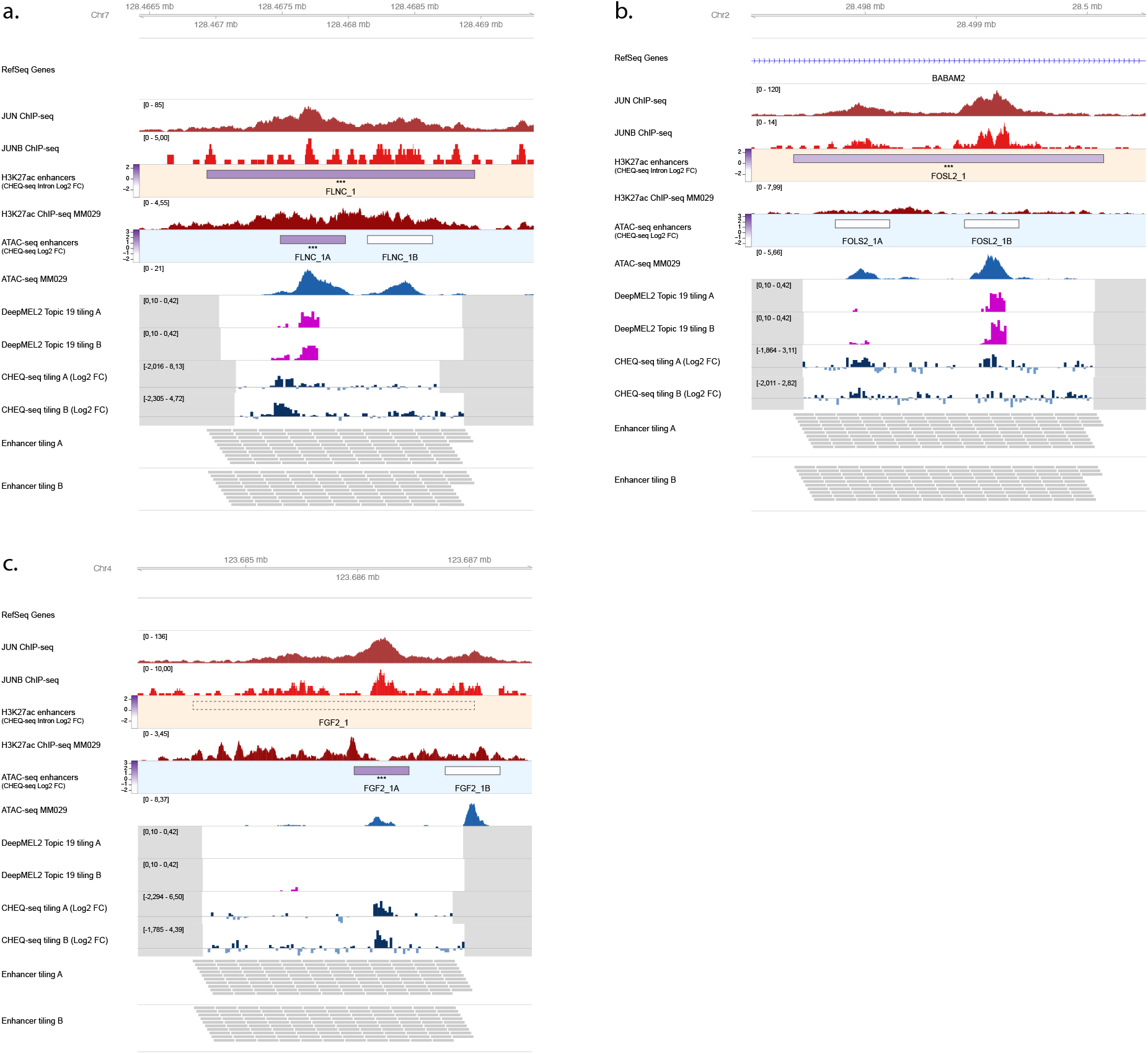
**a.**, Enhancer activity of FLNC_1 (**a.**), FOSL2_1 (**b.**) and FGF2_1 (**c.**) regions. JUN and JUNB ChIP-seq, H3K27ac ChIP-seq and ATAC-seq for MM029 and DeepMEL2 predictions and CHEQ-seq values of the enhancer tiling are displayed. Dark grey areas are regions not covered by the tiling library. CHEQ-seq activity is visible in the **H3K27ac enhancers** and **ATAC-seq enhancers** tracks. Benjamini–Hochberg adjusted p-values: *** < 0.001. Dashed box: regions not recovered following synthesis, cloning or MPRA assay.

**Supplementary figure 6:**
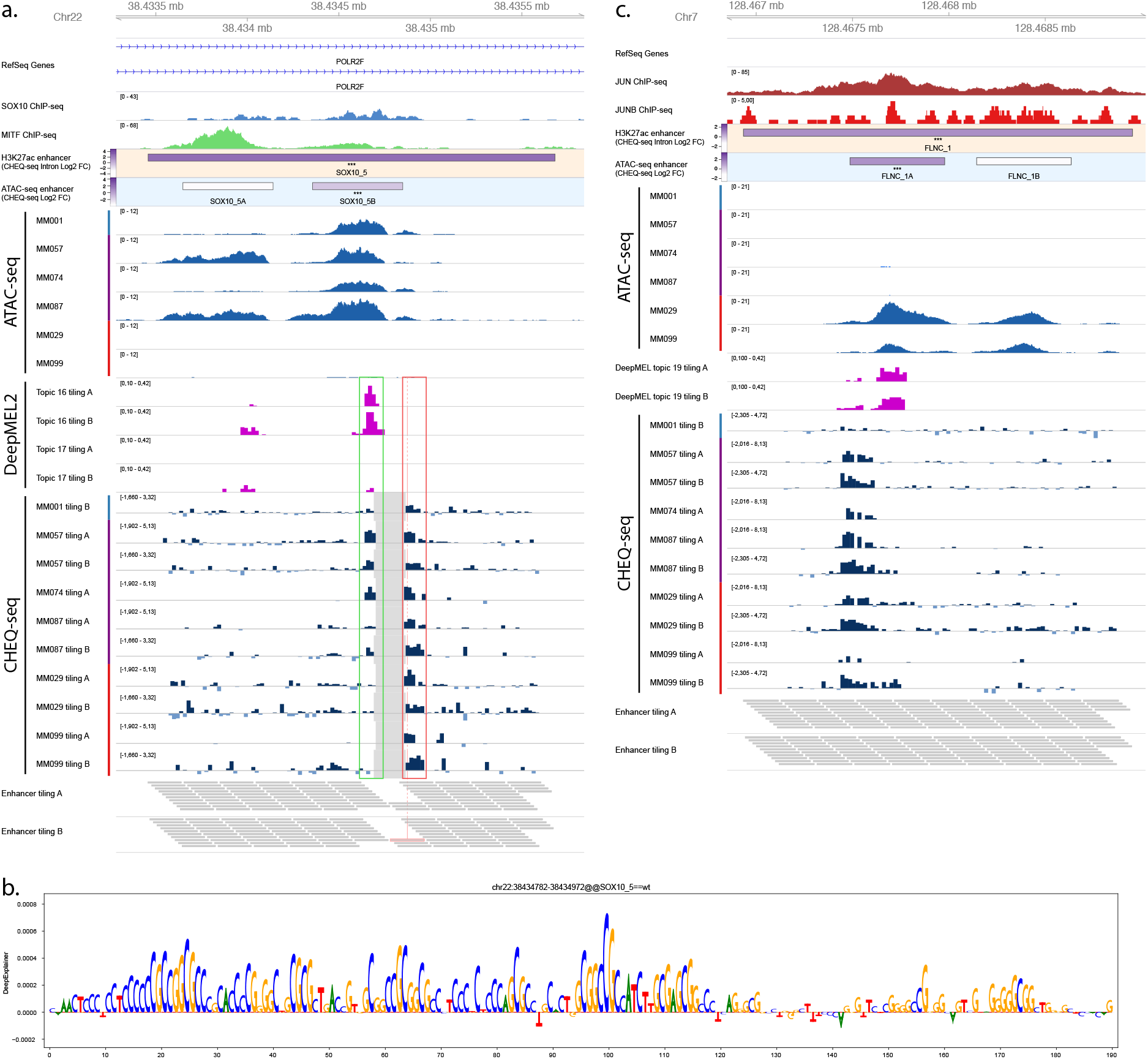
**a.**, Enhancer tiling profile of the SOX10_5 region across all tested MM lines. MM001 activity is shown for the H3K27ac and ATAC-seq regions. The grey area in the CHEQ-seq tiling tracks corresponds to tiles that were not found in the libraries after cloning. The perfect overlap of those missing tiles in libraries A and B suggest that their high GC content caused the synthesis to fail. Bars on the side of the ATAC-seq and CHEQ-seq tracks indicate the phenotype of the cell lines (blue: MEL; purple: MEL intermediate; red: MES). MEL specific and ubiquitous enhancers are highlighted in green and red respectively. **b.**, DeepExplainer profile of the tile highlighted in pink in a. for topic 31 (promoter). **c.**, Enhancer tiling profile of the FLNC_1 region across all tested MM lines. MM029 activity is shown for the H3K27ac and ATAC-seq regions. Benjamini–Hochberg adjusted p-values: *** < 0.001.

**Supplementary figure 7:**
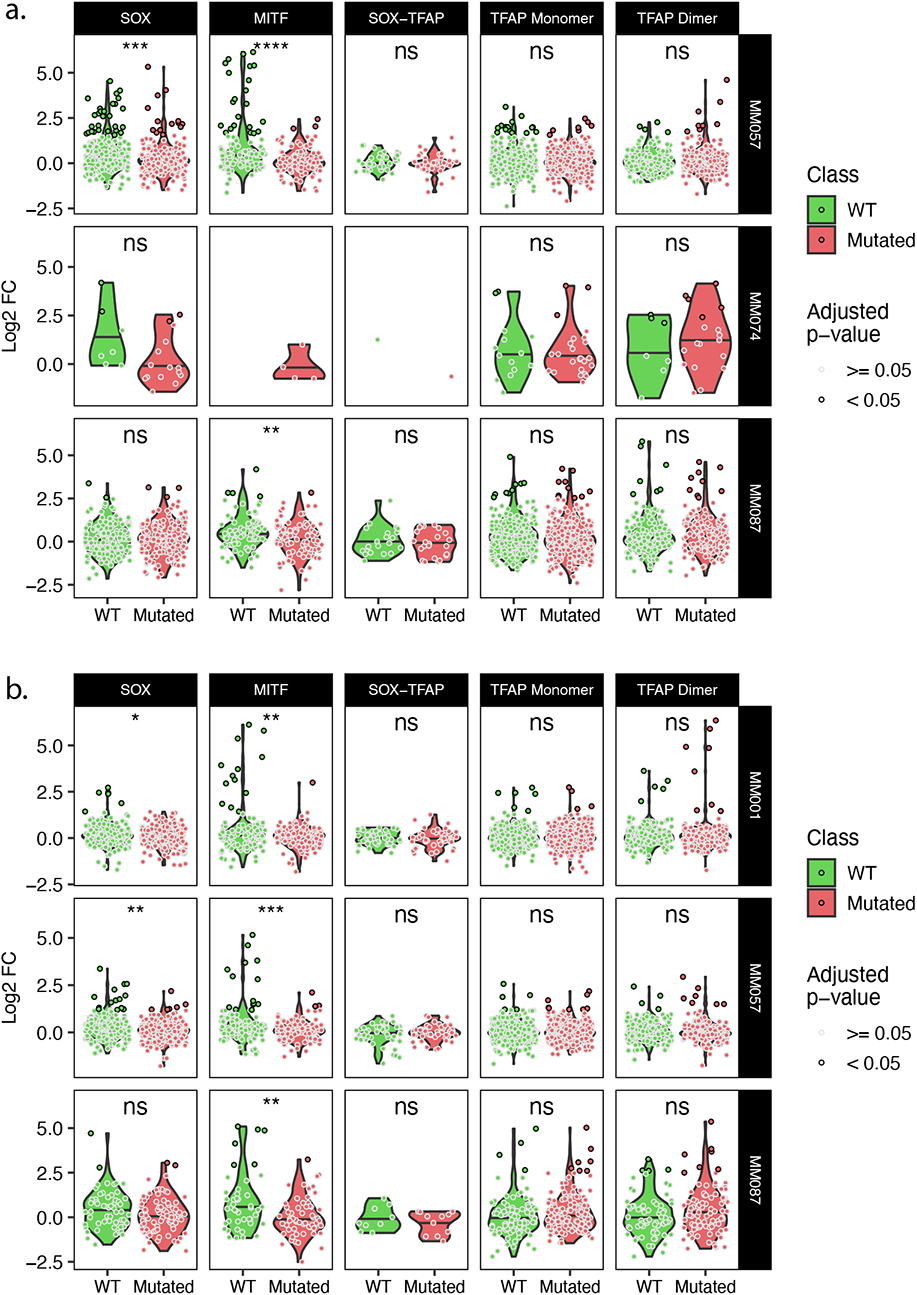
**a.-b.**, Expression of wild type and corresponding mutated tiles for the tiling A (**a.**) and B (**b.**) libraries. T-test p-values: ns > 0.05; * < 0.05; ** < 0.01; *** < 0.001; **** < 0.0001.

**Supplementary figure 8:**
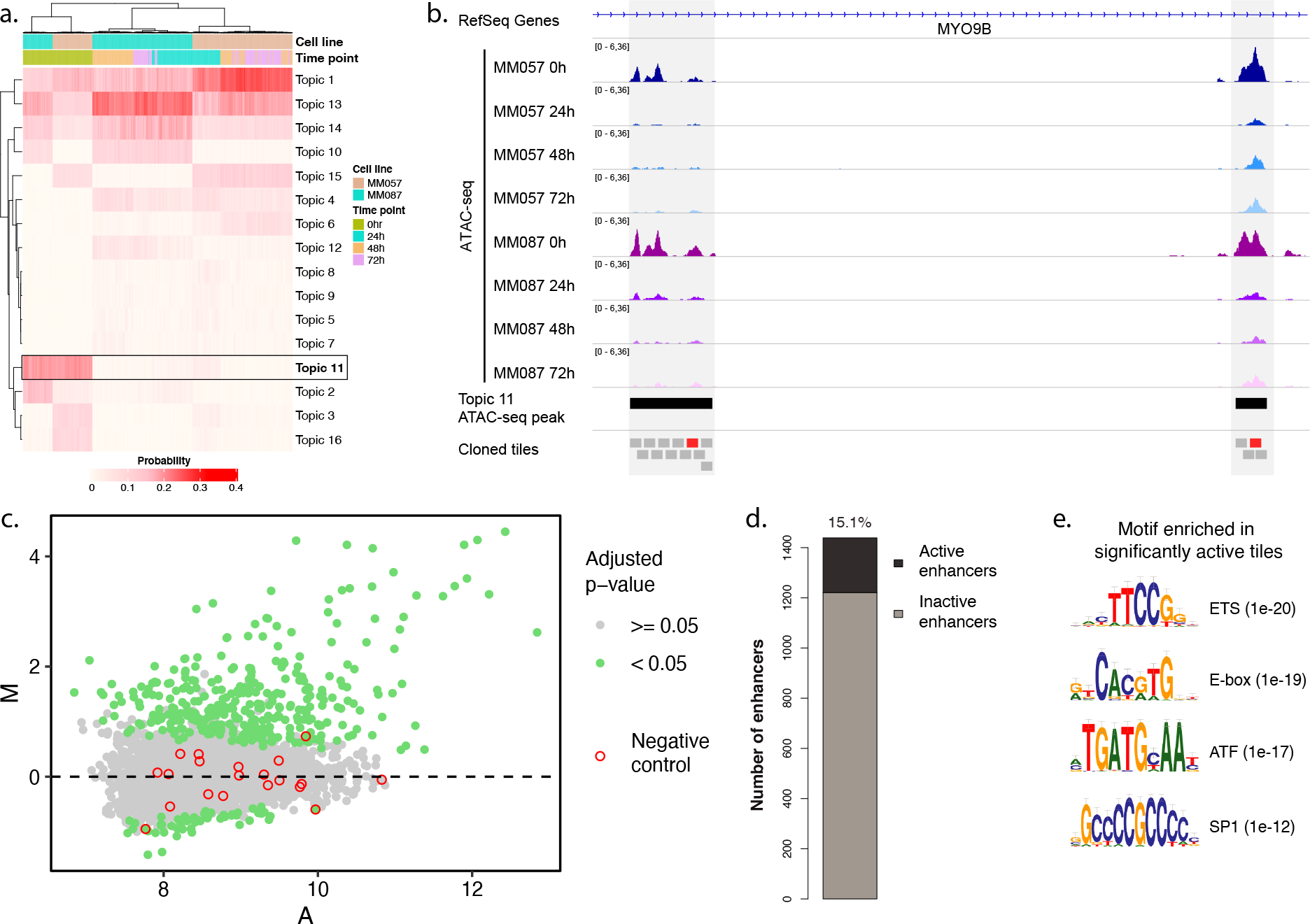
**a.**, Heatmap of topic contribution for each ATAC-seq region over the different cell lines and time points. The highlighted topic 11 contains regions that have reduced predictions following KD. **b.**, ATAC-seq profiles of MM057 and MM087 at 0, 24, 48 and 72h post SOX10-KD. Two topic 11 peaks and 190 bp tiles spanning the enhancers are highlighted by black and grey boxes, respectively. Active tiles are highlighted in red. **c.**, MA plot of all tested tiles in MM087. **d.**, Number of active enhancers. An enhancer is defined as active if at least one of its tiles is active. **e.**, List of the most enriched motifs from the HOMER analysis of active vs inactive enhancers.

**Supplementary figure 9:**
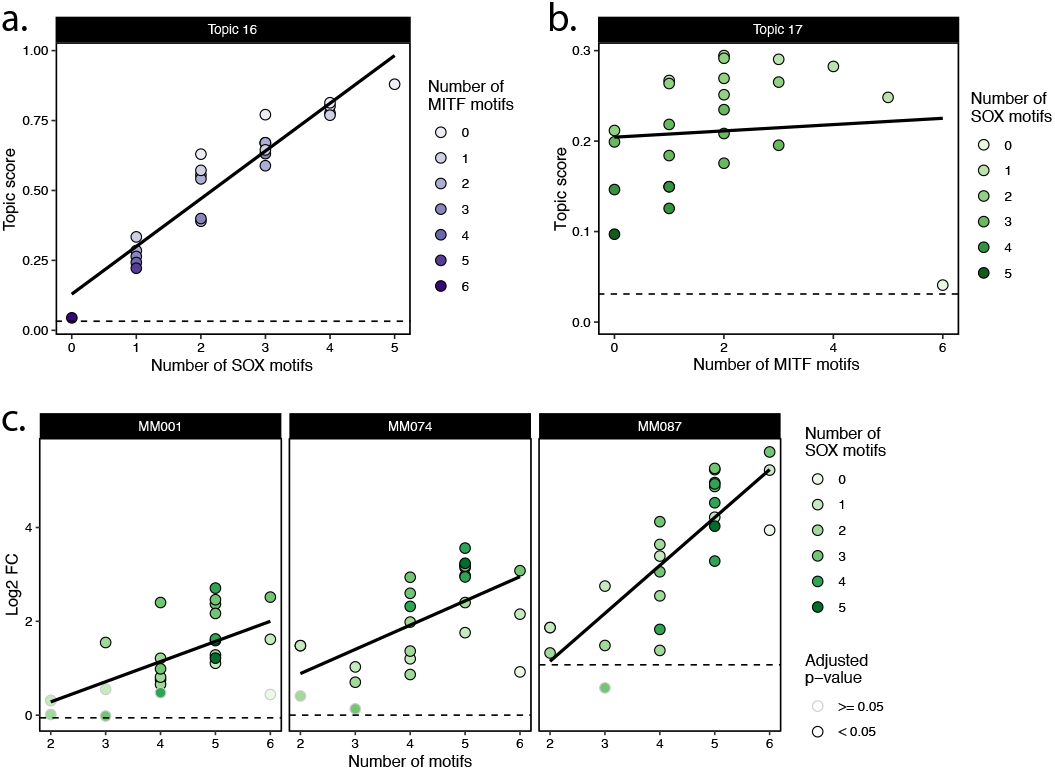
**a.**, Synthetic combinations of SOX and MITF motifs DeepMEL2 prediction scores for Topic 16 with scores ordered by the number of SOX motifs in the sequence. **b.**, Synthetic combinations of SOX and MITF motifs DeepMEL2 prediction scores for Topic 17 with scores ordered by the number of MITF motifs in the sequence. **c.**, CHEQ-seq activity of synthetic enhancers in MM001, MM074 and MM087 sorted by the number of motifs (SOX + MITF) present in the sequence. Dashed line indicates the topic score/log2 FC value of the background sequence without any motif.

**Supplementary figure 10:**
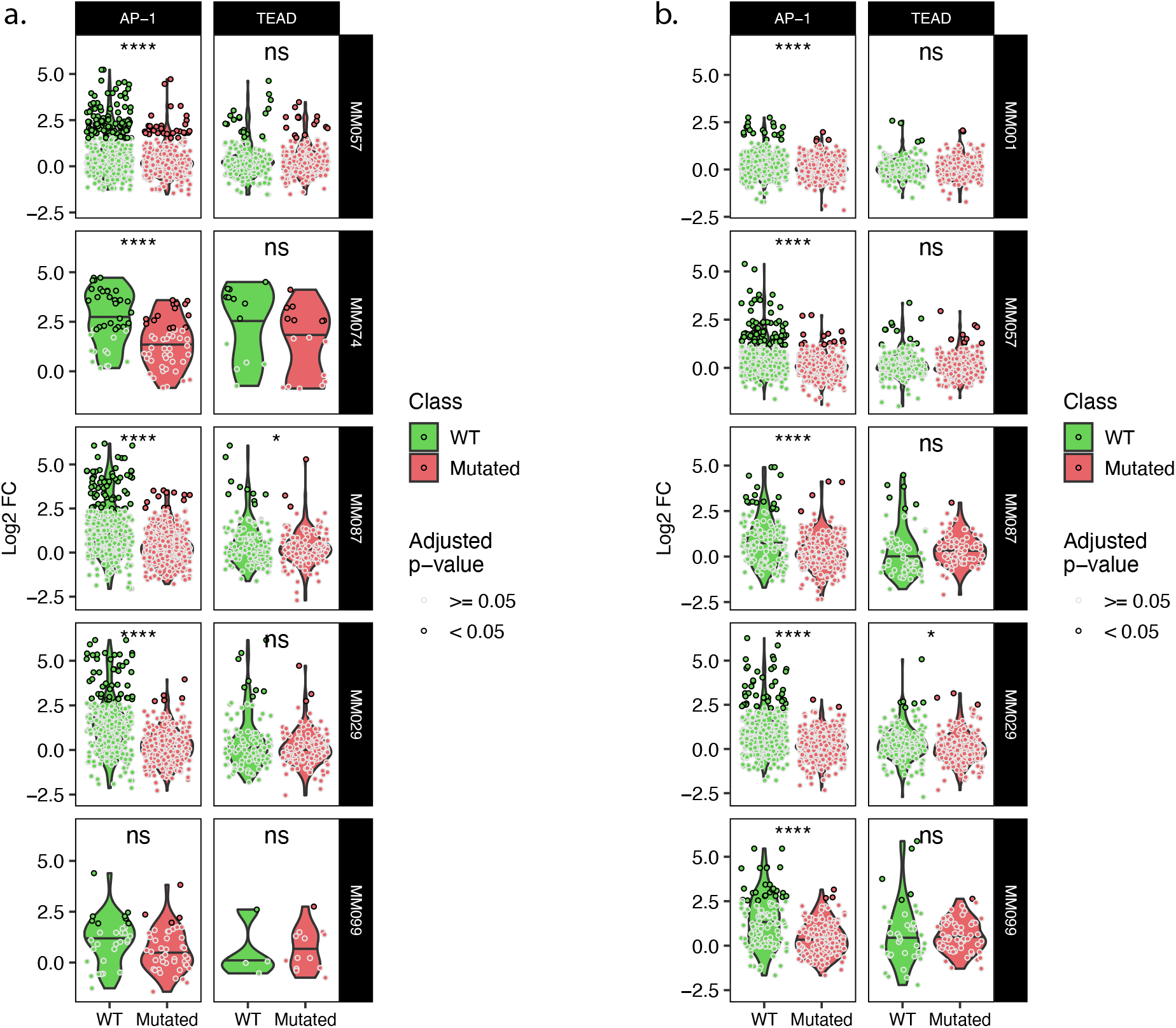
**a.-b.**, Expression of wild type and corresponding mutated tiles for the tiling A (**a.**) and B (**b.**) libraries. T-test p-values: ns > 0.05; * < 0.05; **** < 0.0001.

